# Long-term tillage regime structures bacterial carbon assimilation

**DOI:** 10.1101/2025.01.07.631211

**Authors:** Marie Schaedel, Chantal Koechli, Daniel Buckley

## Abstract

Microbial growth dynamics are deterministic of the fate of carbon in soil, responsible for the transformation of new carbon inputs and their stabilization on soil surfaces. Bacterial life history strategies are predictive of C substrate assimilation and growth response. High disturbance management practices such as tillage alter microbial community structure but have a poorly described impact on life histories that are central to C metabolism. We conducted a DNA stable isotope probing experiment using soil from a long-term field experiment with a 42-year legacy of no-till or annual moldboard plowing. We predicted that divergent legacies of disturbance would result in bacterial communities with distinct life histories, altering C assimilation dynamics. We incubated soil from each tillage regime with ^13^C-xylose and ^13^C-cellulose, two substrates that are components of plant litter and which differ in bioavailability. We identified 730 bacterial taxa that incorporated the labeled substrates and tracked their abundance in bulk microcosm soil over a 30 day period. Carbon addition rapidly altered bacterial community structure and function, with tilled soils demonstrating lower mineralization rates of each substrate. Xylose-assimilating taxa exhibited significantly lagged growth in tilled soils relative to no-till. We also found a higher number and diversity of late (day 30) cellulose incorporators in no-till soil, suggesting that minimal disturbance resulted in a longer residence time of ^13^C-cellulose in members of the bacterial community. We show that soil management practices shape the path of carbon through bacterial communities by altering dynamic growth responses and secondary incorporation of carbon.

**Highlights:**

- DNA SIP identified divergent carbon dynamics resulting from tillage legacies
- Xylose assimilation in plow-till soils was late and decoupled from mineralization
- Cellulose-C was assimilated later in no-till soils relative to plow-till
- Growth responses of incorporator taxa differ by tillage and explain mineralization

## 1. Introduction

Soil microbial communities act as arbiters in determining the fate of carbon (C), with microbial growth response mediating the transformation of C inputs (Sokol et al., 2022) Living microbial communities (bacteria and fungi) facilitate the breakdown of C-containing plant litter into altered biochemical forms (Condron et al., 2010). Such processes are fundamental to the decomposition of plant litter and its transformation into soil organic carbon (SOC; Kallenbach et al., 2016) and stable forms of organic matter (Lavallee et al., 2020). Microbial necromass accounts for an estimated 50-60% of SOC, while living microbes account for a mere 5% of SOC (Camenzind et al., 2023; Sokol et al., 2019). Microbial death and community turnover are therefore fundamental to the accumulation of SOC and its cycling in soil. An increasingly accepted framework proposes that microbial mortality directly influences SOC stocks, as the products of cellular lysis become stabilized on soil surfaces or incorporated in the biomass of another organism (Kästner et al., 2021; Zheng et al., 2021). However, the relationship between microbial growth dynamics and the pathways by which microbial communities transform C in agricultural soils remains unclear.

If microbial necromass comprises a large proportion of SOC, then microbial growth dynamics are central to SOC dynamics. For instance, growth-adapted bacteria that respond rapidly to pulses in nutrient availability tend to exhibit boom-and-bust dynamics, with high growth rates accompanied by high mortality resulting from resource exhaustion or predation (Wattenburger and Buckley, 2023). 16S ribosomal rRNA copy number (*rrn*) has been used as a predictor of growth dynamics and C use efficiency in bacteria (Roller and Schmidt, 2015; Wang et al., 2022). Copiotrophic bacteria contain multiple *rrn* copies in their genomes to rapidly achieve high ribosomal content, enabling them to grow rapidly in response to transient increases in resource availability. Bacteria with different life history strategies and *rrn* copy number exhibit distinct growth responses to C substrates with varying levels of bioavailability (Barnett et al., 2021). For instance, ruderal bacteria rapidly mineralize simple sugars (glucose, xylose), while slower-growing organisms mineralize complex substrates (cellulose, lignin) gradually without rapid variation in population size (Barnett et al., 2021).

While microbial growth and death cycles are deterministic of the fate of C in terrestrial landscapes, we have only recently begun to elucidate linkages between land use, microbial growth dynamics, and C use strategies. Agricultural systems are often characterized by high disturbance management practices such as tillage and fertilization that alter SOC stocks and exert selective pressure on C-cycling microorganisms (Wattenburger and Buckley, 2023; West et al., 2023). Tillage, for instance, mechanically disrupts soil aggregates and accelerates C loss by promoting its rapid oxidation by aerobic microorganisms (Dimassi et al., 2014; Weidhuner et al., 2021). We know from past research that bacterial communities in agricultural soils differ significantly in structure and function from bacterial communities in sites with minimal disturbance, such as forests (Buckley and Schmidt, 2001; Constancias et al., 2014). Compared with undisturbed forest and meadow sites, bacterial communities in agricultural fields experience significant declines in diversity in response to C addition, driven principally by the proliferation of gamma-proteobacteria (Barnett et al., 2022). Furthermore, DNA stable isotope probing (SIP) has revealed that bacteria in agricultural fields preferentially assimilate bioavailable forms of C such as xylose over insoluble forms such as cellulose (Barnett et al., 2022). Accumulating evidence therefore suggests that land use alters bacterial growth dynamics and C cycling, possibly linked to disturbance. Life history frameworks such as Grimes’ competitor-stress tolerator-ruderal (CSR) predict that disturbance disfavors competitors (Grime, 1977).

Bacterial C cycling differs significantly between undisturbed sites and agricultural fields (Barnett et al., 2022), although the influence of specific agricultural management practices on bacterial growth and C cycling is unclear. Agriculture is a significant global source of greenhouse gas emissions, and the ongoing conversion of natural landscapes to agriculture has outsized impacts on SOC loss relative to the area of production gain (Zhang et al., 2021). The total land area dedicated to agriculture accounts for up to 40% of global terrestrial area (Lal, 2004) and it is estimated that agricultural activities to date have released as much as 90 pg C into the atmosphere (Ramankutty et al., 2018). Therefore, improving our understanding of how agricultural practices impact microbial C cycling is necessary to address current trends in SOC loss.

Conservation agriculture practices such as no-till have been associated with SOC sequestration (Bernacchi et al., 2005; Nicoloso and Rice, 2021) and have documented belowground impacts on soil microbial communities (Schmidt et al., 2018; West et al., 2023). Soil microbial diversity has been associated with improved carbon use efficiency, implying a functional connection between community structure and C cycling (Domeignoz-Horta et al., 2020). Although differences in microbial community structure between plow-till and no-till soils are thought to account for differences in C cycling, direct evidence is lacking. Furthermore, the C sequestration potential of no-till practices has been increasingly called into question (Blanco-Canqui, 2021; Powlson et al., 2014) as microbial community responses to such practices tend to be context dependent. Improving our mechanistic understanding of how agricultural management alters microbial C cycling could allow us to better manage for SOC accumulation via microbial pathways.

We conducted a microcosm experiment using ^13^C-xylose and ^13^C-cellulose addition to soils from a long-term field experiment with a 42-year management history of continuous corn cropping with or without moldboard plowing. Both sets of fields are managed with biomass removal, resulting in no-till biomass harvested (NTH) and plow-till biomass harvested (PTH) treatments. Past research at this long-term experimental site has demonstrated significant tillage effects on SOC and organic nitrogen pools (Moebius-Clune et al., 2008). Xylose and cellulose were selected as substrates for this experiment because they are primary components of plant cell walls which differ in their bioavailability due primarily to differences in solubility. The bioavailability of C substrates is closely related to bacterial growth dynamics, with ruderal organisms rapidly metabolizing soluble forms of C and slower-growing competitor species being more active in mineralizing insoluble C (Barnett et al., 2021).

We hypothesized that a long-term tillage regime would exert selective pressure on microbial communities due to disturbance (Grime, 1977). Specifically, we hypothesized that disturbance would alter microbial community structure, causing changes in C mineralization and assimilation dynamics. Furthermore, we predicted that tillage would alter bacterial C assimilation by favoring growth-adapted ruderal taxa that preferentially assimilate soluble substrates such as xylose and disfavoring competitive taxa that preferentially assimilate insoluble substates such as cellulose.

## 2. Methods

### 2.1. Soil sampling

We sampled soils from the long-term tillage research plot at the Miner Institute in Chazy, NY (Clinton County, 44°53.13’N, 73°28.40’W). The research plot is in a factorial 2 x 2 design established in 1973, testing effects of both tillage and residue management. The plot contains four blocks, with each block consisting of four treatments: No-till, returned biomass (NTR); No-till, harvested biomass (NTH); Till, returned biomass (PTR); Till, harvested biomass 122 (PTH). Tillage and biomass treatments were originally established in 1973. Soils for stable isotope probing were sampled in September 2014 from all biomass-harvested (NTH and PTH) plots to focus solely on the effects of tillage. We collected 20 topsoil cores (5 cm depth) across each replicate plot. Cores from a unique replicate plot were combined and homogenized through sieving (2 mm sieve). Soil was placed on ice during transport and stored at 4°C until the beginning of the experiment (3 days). We also collected soil moisture and temperature data for three replicates within each plot.

### 2.2. Microcosm set-up and experimental design

Microcosms were set up in 250 mL Erlenmeyer flasks. 10 g of dry weight soil were placed in the flasks and sealed with butyl rubber stoppers to prevent moisture loss. Dry weight was determined by gravimetric soil moisture measurements for three technical replicates within each tillage treatment and biological replicate (Berthrong et al., 2013). Microcosms were pre-incubated for 2 weeks until the production of CO_2_ stabilized following disruption due to sieving, as assessed by GC-MS (Shimadzu QP2010S GC-MS plumbed with Carboxen-1010 PLOT column, St. Louis, MO) analysis.

Carbon substrates for microcosm enrichment were chosen based on the composition of corn stover (Huang et al., 2009) due to the long-term cropping history of corn in the field site where soils were collected. The substrate solution contained cellulose (0.889 mg C g^-1^ soil), xylose (0.451 mg C g^-1^ soil), arabinose (0.062 mg C g^-1^ soil), mannose (0.034 mg C g^-1^ soil), galactose (0.042 mg C g^-1^ soil), and lignin (0.591 mg C g^-1^ soil). The remaining 18% mass was composed of amino acids (Teknova, #C0705) and Murashige and Skoog basal salt mixture (Millipore Sigma, #M5524). The substrate solution had a C:N ratio of 10, which was comparable to the C:N ratio of the soils used in the experiment. All components of the solution, except for the insoluble C substrates cellulose and lignin, were added in dissolved form at 50% of the water holding capacity of the soil. The two ^13^C treatments substituted 99% ^13^C enriched cellulose or xylose. Both ^13^C and ^12^C cellulose used in the stable isotope probing (SIP) microcosms were prepared using *Gluconoacetobacter xylinus* following Pepe-Renney et al. (2016). The ^12^C control treatments contained carbon with a natural abundance of ^13^C. Water-only control microcosms were treated with an equivalent amount of water and basal salts as other treatments to serve as a control for moisture effects.

Microcosms (n = 112) were prepared using soil derived from field replicates (n = 4). Unlabeled control microcosms and water only controls were destructively sampled five times at 1, 3-, 7-, 14-, and 30-days following substrate addition. Microcosms receiving ^13^C cellulose were sampled on days 3, 7, 14, and 30, while ^13^C xylose microcosms were sampled on days 1, 3, 7, and 14 (Figure 1). Soil from harvested microcosms was stored at –80°C until DNA extraction was performed. Day 30 soils were sub-sampled for isotopic analysis (UC Davis Stable Isotope Facility) and determination of pH, total C, and total nitrogen (N). Soil pH was determined using a 1:1 soil-water slurry method. Total C and N were measured using oven dried, ground samples via a LECO Treu Mac CN-2000 elemental analyzer (LECO Instruments, Lansing, MI) as previously described (Berthrong et al., 2013).

**Figure 1.**
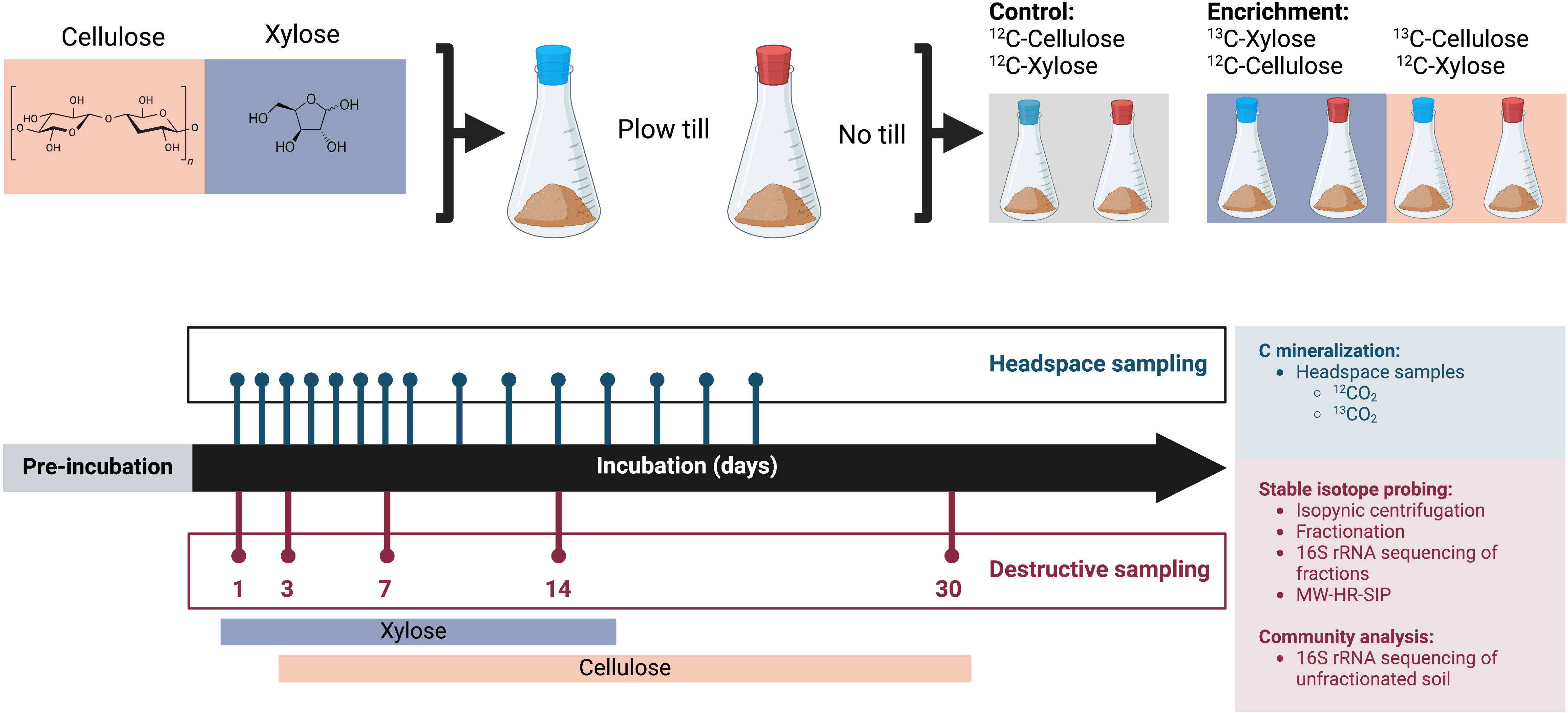

### 2.3. Carbon mineralization

Stoppers were removed from microcosms every two days and the headspace was flushed with filtered (0.2 μm) air. ^12^CO_2_ and ^13^CO_2_ efflux resulting from soil microbial respiration was measured non-destructively at multiple timepoints throughout the 30-day incubation (Figure 1). 2 mL gas sampling vials flushed with helium were used to collect 0.25 mL of microcosm headspace. Headspace samples were then analyzed on a Shimadzu GP2010S GC-MS (St Louis, MO) with an injection port temperature of 200°C. The quadrupole MS was run in selective ion mode (SIM), scanning for m/z of 44 (^12^CO_2_), 45 (^13^CO_2_), and total ion count (TIC). A CO_2_ standard curve was prepared for each sample batch and was used to quantify CO_2_ content in the microcosm headspace.

### 2.4. DNA extraction

We extracted DNA from bulk (unfractionated) microcosms using 2 x 0.25 g of soil from all replicates of each isotope x day x soil combination (112 samples). We used a modified Griffiths phenol-chloroform extraction procedure (Griffiths et al., 2000), as detailed in Supplemental Methods. Following extraction, DNA extracts were cleaned using illustra™MicroSpin™ G-50 columns (GE Healthcare; Buckinghamshire, UK; 27-5330-02) and magnetic bead purification (Agencourt AMPure XP purification; Beckman Coulter; Brea, CA; A63880), according to manufacturer protocols. DNA for isopycnic centrifugation was extracted from 4 technical replicates of 0.25 g of soil, following the phenol-chloroform procedure outlined above. Technical replicates of DNA extractions were pooled and selected for a size of 4 – 14 kb (Youngblut and Buckley, 2014) with a Blue Pippen Prep machine (Sage Science, Beverly, MA) according to the manufacturer’s protocol.

### 2.5. Stable isotope probing and isopycnic centrifugation

Isopycnic centrifugation was performed for a total of 42 samples: all isotope (^12^C vs ^13^C), sampling day, and tillage (NTH vs PTH) treatments from replicate 4 (n=26), and a subset of treatment levels from replicates 2 and 3 (n = 16) which included ^13^C-cellulose microcosms sampled on days 3 and 30 from both tillage treatments. We prepared isopycnic gradients according to a modified protocol based on Neufeld et al. (2007), as described previously (Barnett et al., 2022, 2021). Detailed methods are provided in Supplemental Methods. In brief, size-selected DNA was centrifuged in a CsCl density gradient on an Optima MAX-E ultracentrifuge (Beckman Coulter; Brea, CA). Following centrifugation, 100 µl density fractions were collected and prepared for sequencing (Supplemental Methods).

### 2.6. 16S rRNA amplicon sequencing

We performed amplicon sequencing of the V4 region of 16S rRNA gene across all unfractionated (n = 112) and fractionated (n = 979) samples. We amplified the V4 hypervariable region of 16S rRNA gene using dual-indexed primers (515f / 806r) as described by Kozich et al. (2013). The 16S rRNA libraries were prepared as described previously and in Supplemental Methods. Pooled amplicon libraries were submitted for sequencing at the Cornell Core Facility in Ithaca, NY. Samples were run on an Illumina MiSeq using V2 chemistry with 2 x 250 bp read length.

Raw sequence data were processed using QIIME2 v2023.9.1 (Boylen et al, 2019). Fractionated samples from SIP and unfractionated microcosm samples were processed concurrently in the same pipeline to enable amplicon sequence variant (ASV) cross-referencing between SIP fractions and unfractionated microcosm samples. As described previously (Wattenburger and Buckley, 2023), sequences were demultiplexed and trimmed based on an examination of the quality scores. Quality filtering, denoising, and ASV merging was performed with DADA2 v2023.9.1. Phylogenetic assignment of ASVs was performed using the Silva v138 database (Quast et al., 2013). We used MAFFT 7 (Katoh and Standley, 2013) to align sequences and FastTree 2 (Price et al., 2010) to construct a phylogenetic tree. Processed sequences from fractionated and unfractionated samples were deposited in the NCBI Short Read Archives (BioProject PRJNA1170979). All scripts used in processing and analyzing the sequence data can be accessed at https://github.com/schaedem/chazy_sip.

Processed sequences were imported into R version 4.0.3 (R Core Team, 2020). ASV count data from unfractionated samples were rarefied to an even depth of 1,567 in phyloseq version 1.34.0 (McMurdie and Holmes, 2013) using the *rarefy_even_depth* function. Alpha diversity indices were calculated using *estimate_richness* in phyloseq. Pielou’s evenness was calculated as the ratio of Shannon diversity to the natural log of richness per sample. Weighted and unweighted Unifrac distances were calculated with the *Unifrac* fuction in phyloseq, and Bray-Curtis distances were generated with *avgdist* in vegan (version 2.5.7; Oksanen et al., 2022). The estimated 16S rRNA copy number (*rrn*) for each ASV was determined using paprica (Bowman and Ducklow, 2015). Faith’s phylogenetic diversity was determined using *pd* from the picante package (Kembel et al., 2010).

### 2.7. Identification of ^13^C-labeled taxa

MW-HR-SIP was performed using the HTSSIP package (Youngblut et al., 2018) in R. ASV count data was pruned in phyloseq using the *filter_taxa* function prior to identifying incorporators to retain ASVs that occurred at least twice across all samples. We compared the distribution of taxa within overlapping density windows in the range of 1.70 to 1.77 g ml^-1^ (Youngblut et al., 2018) and used DESeq2 (Love et al., 2014) to identify differentially abundant ASVs in ^13^C labeled treatments. ASVs were deemed ^13^C incorporators if they exhibited a log_2_-fold enrichment of at least 0.25 in heavy isotope fractions with a Benjamini-Hochberg corrected p-value of <0.05 relative to ^12^C fractions. The number of comparisons was minimized by performing independent sparsity filtering at each overlapping density window. We used venn diagrams to visualize shared incorporators by tillage regime and substrate with the eulerr package in R (Larsson, 2024).

### 2.8. Statistical analysis

#### 2.8.1. Carbon mineralization

We evaluated the contribution of tillage history and days since C addition on cumulative mineralization and mineralization rates of individual substrates (^13^C) and combined substrates (^12^C+^13^C) within the microcosm headspace. All analyses were performed in R version 4.0.3 (R Core Team, 2020), and data frame manipulation was performed with tidyverse functions (Wickham et al., 2019). We used linear mixed effects models from the lme4 package (Bates et al., 2015), specifying sample day and land use as fixed effects and replicate as a random effect. The significance of individual and interactive terms within the models were assessed with *anova* from the stats package (R Core Team, 2020). To identify significant contrasts in mineralization between tillage histories, we used *lsmeans* from the emmeans package (Lenth et al., 2022) and corrected p-values for multiple comparisons with the Benjamini-Hochberg adjustment. Finally, we used *t.test* in the stats package to assess endpoint differences in cumulative ^13^C mineralized for each substrate and in total for all substrates (^12^C and ^13^C) by tillage regime. The chosen endpoints for each substrate were day 14 for xylose and day 22 for cellulose. Carbon mineralization data were normally distributed according to Shapiro-Wilk tests.

#### 2.8.2. Microcosm community composition

We investigated the change in community composition following C addition in unfractionated microcosm samples by computing alpha and beta diversity at each timepoint. The change in diversity was computed as the difference between microcosms receiving a C substrate solution (^12^C + ^13^C) and the baseline water-only control that did not receive carbon. We employed linear mixed effects models as described previously to identify the contribution of tillage, sampling day, and their interaction in driving changes to diversity. Significance was determined using Benjamini-Hochberg corrected p-values of less than 0.05. Homogeneity of dispersion across tillage regimes was determined using *betadisper* and tested for significance with *permutest* from vegan. We identified differentially abundant taxa using a Maaslin2 (Mallick et al., 2021) model in which tillage, sampling day, and their interaction were specified as fixed effects. Microcosm replicate (corresponding to the field replicate from which soils were collected) was additionally specified as a random effect.

#### 2.8.3. Growth responses and life history traits of incorporator taxa

Maximum log2-foldchange (max l2fc) for incorporator ASVs was determined using normalized abundance values for each ASV in bulk microcosm soil. Following Barnett et al. (2021), ASV relative abundance was divided first by predicted *rrn* and then by the estimated *rrn* of the entire sample. These normalized abundances were weighted by DNA yield (mg DNA g^-1^ soil), which was measured with the Quant-iT PicoGreen dsDNA Assay Kit (Termo Fisher Scientific). Max l2fc was calculated for each ASV as the difference between baseline abundance in untreated control soil (receiving H_2_O and micronutrients but no carbon) and the maximum recorded abundance after C addition. If the maximum normalized abundance of an incorporator in treated soil was less than its baseline abundance in untreated soil, max l2fc for that incorporator was assigned a value of zero. We defined latency as the difference between when ^13^C assimilation was detected in an incorporator ASV and the day of peak mineralization for a given substrate. For calculating latency, peak mineralization for xylose was assigned day 2, while cellulose was assigned day 6. ASVs that had a detected ^13^C label before the time of peak mineralization were assigned a latency value of zero. The degree of ^13^C assimilation was assigned for each incorporator ASV as the log2-fold change in abundance between ^12^C and ^13^C SIP density fractions (Wilhelm et al., 2021).

We used Wilcoxon rank-sum tests for 2-factor comparisons evaluating incorporator traits according to substrate and tillage (*wilcox.test* in the stats package; R Core Team, 2020). Wilcoxon tests were used to evaluate differences in predicted *rrn,* latency, species richness, and Faith’s phylogenetic diversity (calculated using *pd* from picante; Kembel et al., 2010) among incorporators from divergent tillage regimes. Relationships between *rrn* and growth traits (max l2fc, degree of ^13^C assimilation, and latency) were assessed with Spearman rank correlation as described previously.

Lastly, we assessed relationships between C cycling dynamics and phylogeny using functional distance matrices for incorporator ASVs in PTH and NTH microcosms. Carbon substrate (xylose or cellulose), the day of label detection, and the degree of enrichment (log_2_-fold enrichment in heavy fractions) were considered functional responses relevant to C metabolism. These values were scaled using *decostand* and functional distance was calculated using *vegdist(method = “altGower”)* in vegan. Phylogenetic distance between incorporator ASVs was determined using *cophenetic.phylo* in phyloseq (McMurdie and Holmes, 2013). We evaluated the relationship between phylogenetic and functional distance among incorporator taxa for each tillage regime separately using *cor.test(method = “spearman”)* from the stats package in R (R Core Team, 2020).

## 3. Results

### 3.1. Mineralization dynamics of xylose and cellulose

Equal amounts of cellulose and xylose were added to soil microcosms and CO_2_ production (as ^12^C + ^13^C) was tracked over time. Cumulative total C mineralization rate (^12^C + ^13^C) differed with respect to tillage history (mixed effects ANOVA, *p* < 0.001), time (mixed effects ANOVA, *p* < 0.001), and the interaction between tillage and time (mixed effects ANOVA, *p* < 0.001; Supplemental Table 2). Cumulative C mineralization rates differed with respect to tillage on days 1 and 2, with no-till (NTH) soils exhibiting higher total mineralization rates relative to plow-till (PTH) soils (Supplemental Figure 1). NTH microcosms exhibited early differences in the cumulative amount of xylose mineralized (mg C), while tillage regimes did not differ in cumulative mineralized cellulose until day 7 (Figure 2). By the sampling endpoints (day 14 for xylose, day 22 for cellulose), NTH communities mineralized more ^13^C-cellulose than PTH communities (t-test, *p* < 0.04) but there was no difference between tillage regimes in the amount of xylose mineralized (Supplemental Figure 2, Supplemental Table 3). NTH had higher rates of ^13^C-cellulose mineralization than PTH on days 6 through 10 and higher rates of ^13^C-xylose mineralization on day 2 (Figure 3). These results indicate that tillage history impacted mineralization dynamics of the two C substrates, especially in the early period of xylose mineralization and throughout the incubation period for cellulose.

**Figure 2.**
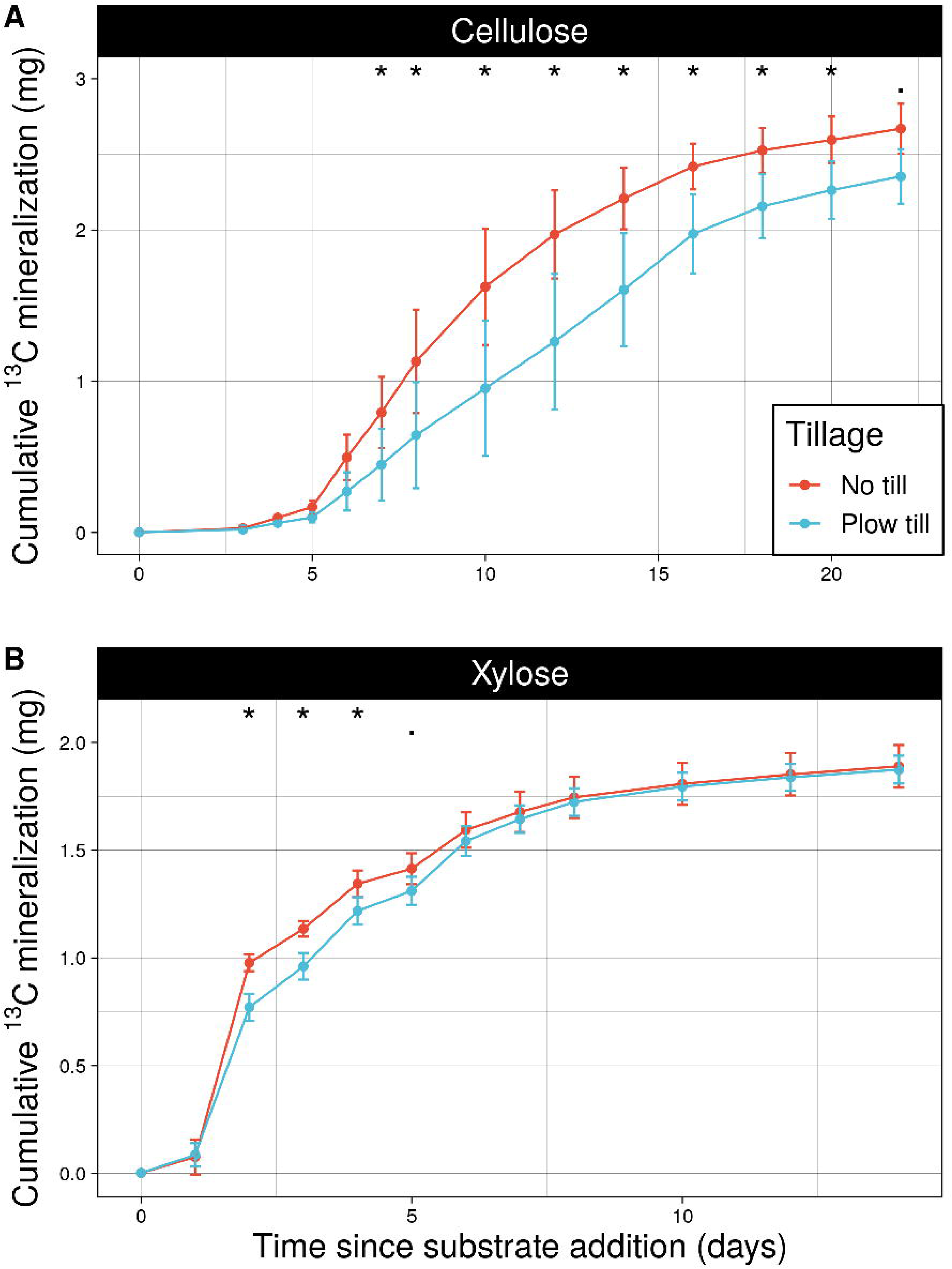

**Figure 3.**
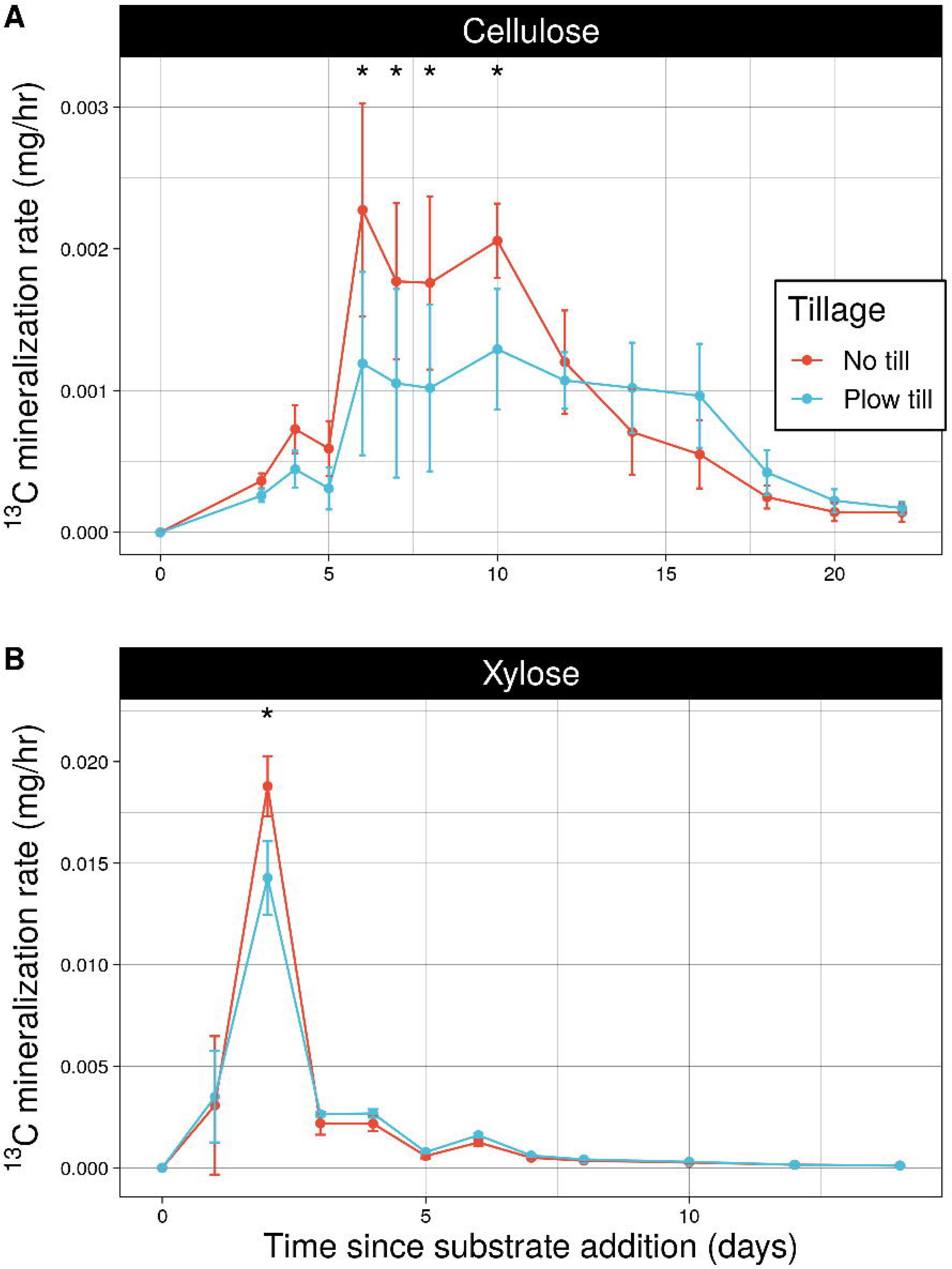

### 3.2. Tillage alters bacterial community responses to carbon

We assessed bacterial community composition in microcosm bulk soil and tested for differences between tillage regimes. Bacterial community membership in the microcosms was dominated by ASVs assigned to *Proteobacteria* and *Actinobacteria* (Figure 4A). NTH soils exhibited greater evenness than PTH soils (mixed effects ANOVA, *p* < 0.01; Supplemental Table 4), but no differences in richness, Shannon diversity, or Inverse Simpson diversity were observed between tillage regimes (Supplemental Table 4). We identified a total of 89 ASVs that were differentially abundant (Maaslin2) with respect to tillage history after accounting for sampling day. Differentially abundant taxa had uniformly higher abundance in PTH soil and belonged exclusively to the phyla *Actinobacteria, Bacteriodota,* or *Proteobacteria*. DNA yield (ng ul^-1^), which we measured as a proxy for microbial biomass (Barnett et al., 2022; Fornasier et al., 2014), did not correlate with evenness, nor did it differ between PTH and NTH soil.

**Figure 4.**
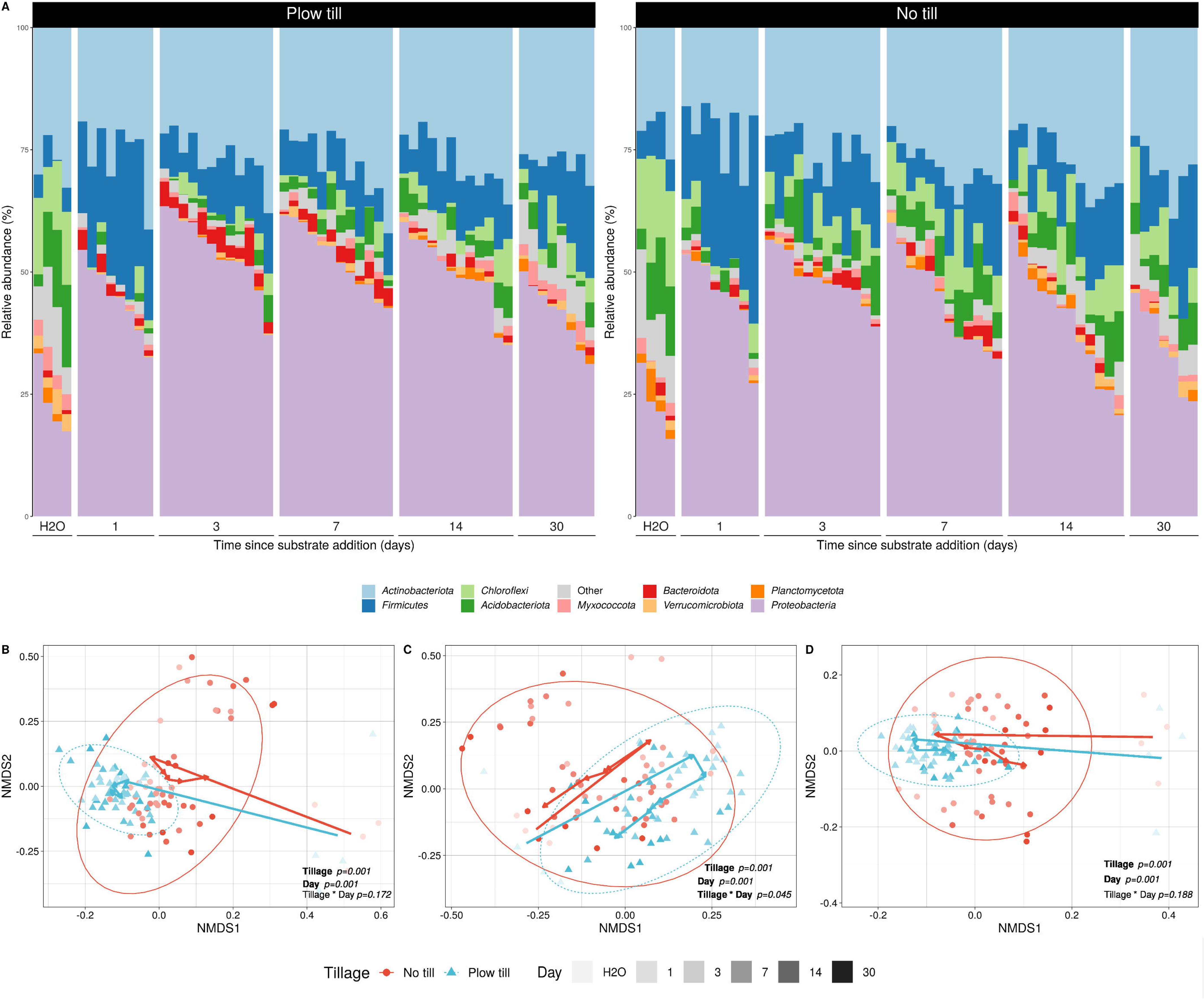

We observed differential responses to C addition between the two tillage regimes. The degree to which C addition altered Bray-Curtis and weighted UniFrac distances over time was similar across tillage regimes: communities that received C inputs remained significantly different from water-only controls throughout the 30-day incubation following C addition (Figure 4B,D; Supplemental Fig 5A,B). In contrast, the unweighted UniFrac distance, which does not take taxon abundance into account, resembled untreated control communities after an initial change in community structure immediately following C addition. The unweighted Unifrac distance of NTH communities differed from the baseline for the first 7 days following C addition while PTH communities differed from the baseline for 14 days (Figure 4C; Supplemental Fig 5C). Taken together, these results suggest that C addition altered community structure and evenness by impacting taxon abundance. We also observed significantly more dispersion in NTH communities than PTH communities throughout the incubation (Fig 4B-D; PERMDISP p<0.001 for weighted Unifrac and Bray-Curtis).

### 3.3. Bacterial carbon assimilation differs by tillage regime

To probe differences in bacterial C assimilation across tillage regimes, we employed multiple-window high-resolution DNA-SIP (MW-HR-SIP; Youngblut et al., 2018). Amplicon sequence variants (ASVs) that incorporated ^13^C into their DNA from either ^13^C-xylose or ^13^C-cellulose were identified based on significant shifts in buoyant density relative to microcosms receiving ^12^C carbon. We identified 730 unique ASVs that assimilated ^13^C into their genomes. Of these, 437 ASVs were labeled in NTH microcosms, while 468 were labeled in PTH microcosms. A total of 538 incorporator taxa were detected in both soils, but the labelling pattern for most taxa differed with respect to tillage history (Fig 5). We found a similar number of ^13^C-cellulose incorporators in both soils (372 in NTH, 352 in PTH), but approximately 2.4 times more unique taxa incorporated ^13^C-xylose in PTH soils relative to NTH soils (234 in PTH and 96 in NTH). Only 337 of the 730 incorporator ASVs were detected in the unfractionated soil microcosm communities. DNA-SIP can detect rare taxa that often go undetected in standard DNA sequencing approaches; hence, many taxa (53.8%) participating in C cycling were present at very low abundance.

**Figure 5.**
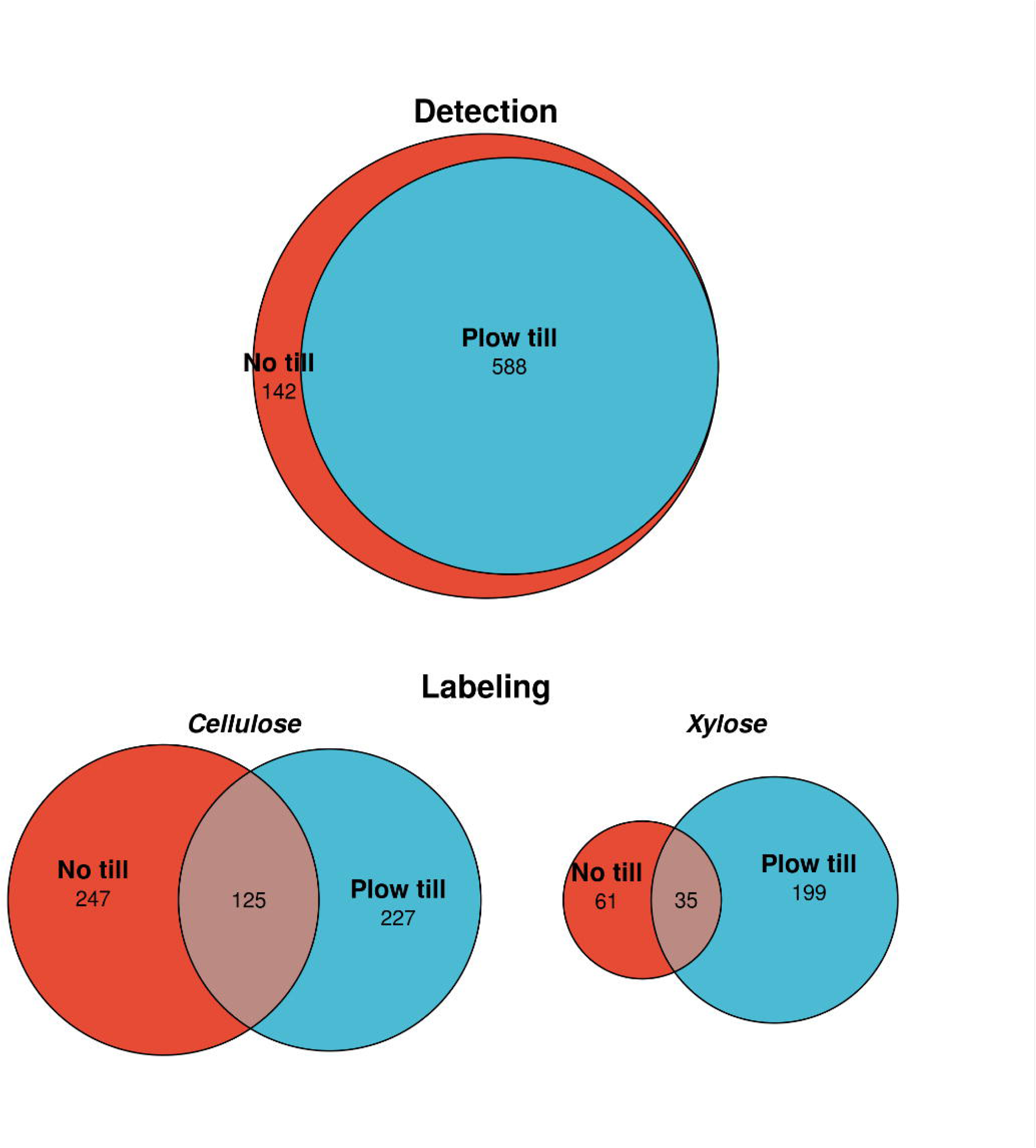

Incorporator diversity (Faith’s phylogenetic diversity and species richness) varied with respect to tillage, days since C addition, and their interaction (Supplemental Table 6). Early NTH xylose incorporators demonstrated higher diversity immediately after substrate addition on day 1 (Supplemental Figure 7). PTH xylose incorporators were most diverse by day 7, coinciding with higher abundance and dual labeling with cellulose (Figure 6A, 6B). Late cellulose incorporators were more diverse in NTH than PTH microcosms by day 30 (Supplemental Figure 7). Temporal patterns in incorporator diversity largely mirrored detection of labeling and normalized abundances of incorporators in microcosm communities (Fig 6A).

**Figure 6.**
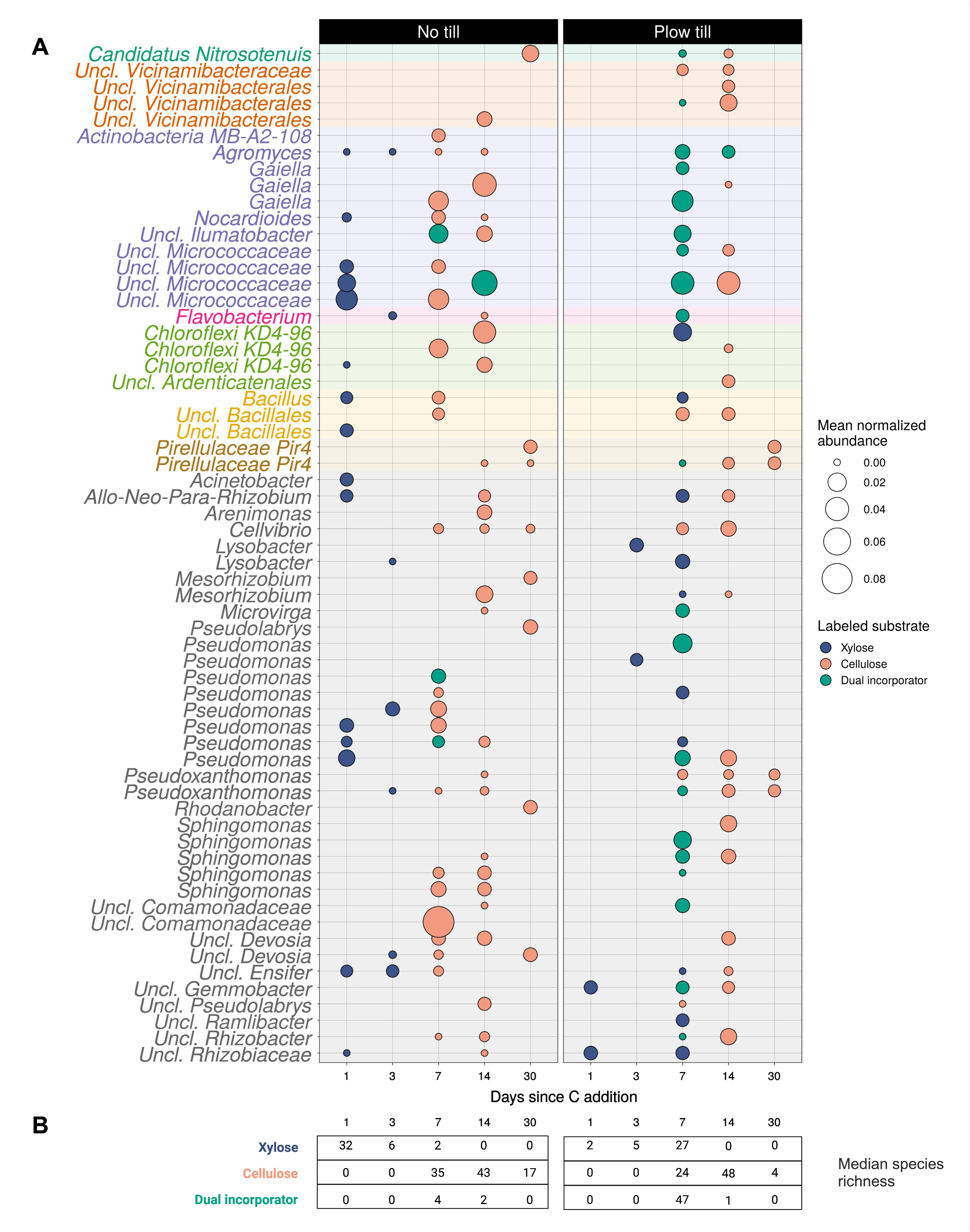

### 3.4. Incorporator growth dynamics differ by tillage and carbon source

We hypothesized that 42 years of tillage would alter C assimilation and growth dynamics due to selective pressures associated with disturbance (Grime, 1977). We calculated the predicted ribosomal RNA copy number (*rrn*) of each incorporator ASV due to its value as an indicator of life history, predictive of lag time (Stevenson and Schmidt, 2004) and translational power (Roller et al., 2016). Xylose incorporators had a median *rrn* of 5, while cellulose incorporators had a median *rrn* of 3 (Wilcoxon rank-sum test, p < 0.001). Cellulose incorporator *rrn* did not differ by tillage regime, although xylose incorporator *rrn* was higher in NTH (median *rrn* of 5) than in PTH (median *rrn* of 2.6; Wilcoxon rank-sum test, p < 0.001; Figure 7A).

**Figure 7.**
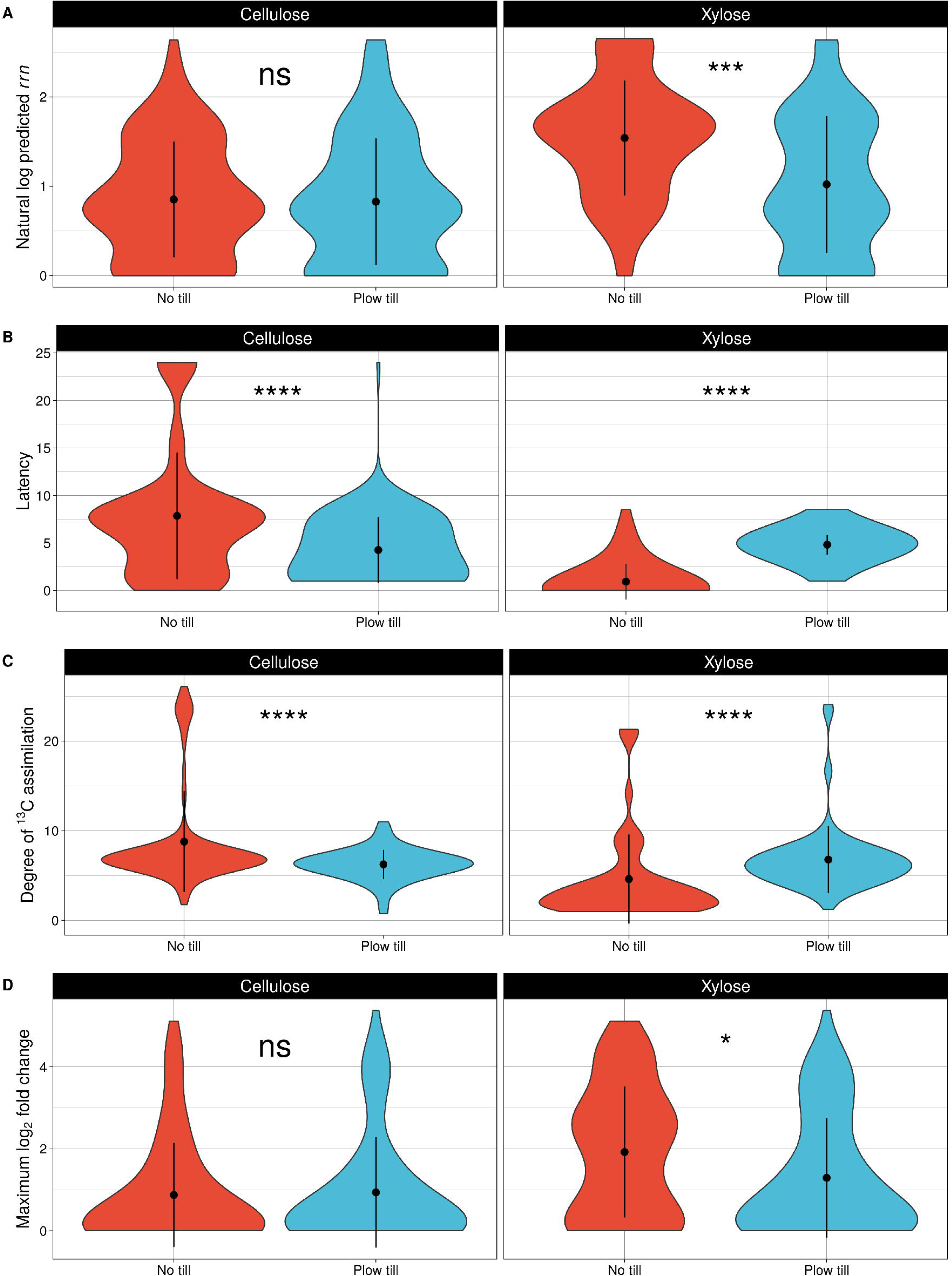

Furthermore, C assimilation and growth dynamics differed among incorporators according to tillage regime, *rrn*, and substrate. Xylose incorporators exhibited an earlier growth response in NTH than PTH. Xylose labeling in NTH soils occurred mostly (69.6%) on day 1 and was dominated by high *rrn* ASVs (Fig 6, Supplemental Fig 7). In PTH soils, 90.2% of xylose incorporation occurred late on day 7 outside of peak mineralization activity and was dominated by a diverse mixture of high and low *rrn* ASVs. In contrast, cellulose incorporators in NTH had a later growth response than PTH cellulose incorporators. More cellulose incorporators were labeled early on day 7 in PTH (57.3%), then in NTH (38.6%; Figure 6, Supplemental Fig 7). Long-term tillage therefore resulted in a lagged growth response of high *rrn* xylose assimilators as well as the dual labeling of cellulose and xylose by low and high *rrn* taxa on day 7.

Incorporator latency, the difference between detection and peak mineralization activity, also differed by C substrate and tillage (Wilcoxon rank sum test, p<0.001; Fig 7B). Cellulose incorporators had higher median latency in NTH (8 days) than in PTH (4 days). In contrast, xylose incorporators had higher latency in PTH (5 days) than in NTH (0 days). Thus, disturbance through tillage altered growth responses according to *rrn* and C substrate, resulting in lagged xylose assimilation. We also examined maximum log2-foldchange (max l2fc), which is an indicator of dynamic growth potential and has been associated with ruderal life history strategies (Barnett et al, 2023). As expected, max l2fc was positively associated with *rrn* among incorporators from both tillage histories (Supplemental Figure 8). Cellulose incorporators did not differ by max l2fc according to tillage regime, although the median max l2fc of xylose incorporators was 3-fold higher in NTH than in PTH (Wilcoxon rank-sum test, p = 0.03; Figure 7D). Our results confirm an expected relationship between *rrn* and max l2fc, and indicate a higher potential for dynamic growth among xylose incorporators in NTH relative to PTH.

Next, we examined the degree of ^13^C labeling among incorporators across the two tillage regimes and C substrates (see Methods). The degree of labeling was negatively associated with incorporator *rrn* in NTH soils but was only weakly associated with *rrn* in PTH soils (Supplemental Fig 8). NTH cellulose incorporators had a significantly higher degree of labeling than PTH cellulose incorporators (Wilcoxon rank-sum test, p<0.001). Conversely, PTH xylose incorporators had a higher degree of labeling than NTH (Wilcoxon rank-sum test, p < 0.001; Fig 7C). Differences in the degree of ^13^C assimilation suggest that incorporators from divergent tillage regimes differed in sources of primary labeling.

Finally, we examined whether growth responses to C (latency, max l2fc, degree of ^13^C assimilation) were phylogenetically conserved with respect to tillage. C cycling responses exhibited little phylogenetic conservation beyond the genus level regardless of tillage regime (Supplemental Figure 8). Growth responses to C among incorporators were more strongly related to phylogenetic distance in NTH soils (Spearman’s *ρ* = 0.17, p<0.001) than in PTH soils (Spearman’s *ρ* = -0.07, *p* = 0.001).

## 4. Discussion

### 4.1. Tillage altered carbon mineralization rates and total carbon mineralization

We found that a 42-year history of moldboard plowing significantly altered microbial mineralization dynamics of xylose and cellulose, two carbon substrates that are major components of plant cell walls and which differ in bioavailability. We show that NTH microcosms mineralized more xylose than PTH microcosms within 5 days of substrate addition. NTH microcosms also mineralized more cellulose than PTH microcosms throughout the experiment (Supplemental Figure 2). Thus, our findings suggest faster and more complete microbial processing of carbon in no-till soils relative to tilled soils.

Our results highlight the important role of historical disturbance in contributing to divergent C dynamics in managed ecosystems. Our findings agree with previous reports that link long-term management with the metabolic potential of microbial communities. For instance, Barnett et al. (2022) observed significantly delayed C mineralization in cropland soils compared to forest and undisturbed meadow sites. Both soils used in the present study were managed with residue removal for 42 years, with tillage as the sole variable, resulting in significantly higher organic matter in NTH soils relative to PTH soils (Moebius-Clune et al., 2008; Supplemental Table 1). Therefore, the differences in C mineralization and incorporation we observed are likely linked to microbial growth dynamics that underlie the legacy effects of tillage (ie, disturbance) on soil C cycling.

### 4.2. Tillage structured bacterial community responses to carbon

We show that tillage legacy altered bacterial community structure and response to carbon. Beta-diversity (Bray-Curtis and weighted/unweighted Unifrac) varied by tillage history and time since C addition. Carbon input to PTH was accompanied by dynamic increases in the relative abundance of *Proteobacteria, Actinobacteria,* and *Bacteroidota* (Fig 4A). Both *Proteobacteria* (syn *Pseudomonodata*) and *Actinobacteria* have been previously linked to legacies of disturbance (Mickan et al., 2019; Seitz et al., 2021) or artificially imposed disturbance (West and Whitman, 2022). We also found that many taxa within *Proteobacteria*, including those belonging to the genera *Pseudomonas* and *Rhizobiaceae,* were highly abundant rapid responders to C addition in both tillage regimes (Fig 6A). Thus, tillage resulted in divergent responses to C within bacterial communities, likely related to the proliferation of disturbance-adapted taxa in PTH microcosms.

Plow-till microcosm communities were also less dispersed than no-till communities. This observation concurs with prior literature that documents the homogenizing influence of disturbance on microbial communities (West and Whitman, 2022). We interpret our results as providing further evidence for the significant contribution of land management to bacterial community assembly. By inverting topsoil and disrupting soil physical structure, tillage may structure bacterial communities through homogenizing dispersal (West et al., 2023) and reducing microsite variation in community diversity associated that are with niche habitats in undisturbed soil aggregates (Dong et al., 2021; Mummey et al., 2006; Nunan et al., 2003). Given prior evidence, we expect that stochastic processes dominate bacterial community assembly in tilled soil (Li et al., 2021, 2023; West et al., 2023). The impact of community assembly processes on carbon cycling has not previously been addressed, yet our results suggest that the more homogenous communities associated with tilled soils exhibited heightened susceptibility to further disturbance in the form of C addition. It is unclear how stochasticity alters the C cycling function of bacterial communities. However, it is evident that the legacy impacts of tillage structured community-wide responses to C addition that were not explained by overall differences in diversity or initial community size (assessed by DNA yield). If microbial biomass and diversity are lower in arable soils relative to other land uses (Lee et al., 2020; Nsabimana et al., 2004; Zhang et al., 2016), community assembly processes may have an outsized influence on carbon assimilation dynamics by influencing the successional state of the community.

### 4.3. Tillage differentially altered cellulose and xylose assimilation processes

Our results indicate that the historical disturbance effect of tillage resulted in fundamental differences in bacterial growth dynamics and carbon mineralization. In line with our hypothesis that tillage would favor the proliferation of pulse-adapted organisms, we identified a greater number of xylose incorporators in PTH soils than in NTH soils. However, we also found that 90.2% of xylose-incorporating bacterial ASVs in PTH microcosms were labeled on day 7, decoupled from peak xylose mineralization that occurred on day 2. This suggests that high *rrn,* ruderal organisms started at lower abundance in PTH soil relative to NTH. PTH xylose incorporators therefore required a longer lead time after being primed by C addition to reach their maximal abundance (Supplemental Figure 7). Xylose and cellulose were consequently assimilated by many of the same ASVs in PTH, regardless of *rrn* copy number. In contrast, there was no lag in xylose incorporation in NTH soil, evidenced by significantly lower latency of labeling and a majority of high *rrn* incorporators reaching peak abundance within one day following C addition. This finding concurs with Barnett et al. (2022)’s observation of lagged xylose mineralization in disturbed cropland soil relative to undisturbed meadow and forest soils. Importantly, it suggests that minimal disturbance management practices such as no-till can shift microbial carbon metabolism to more closely resemble the functions of undisturbed soil.

Earlier labeling in no-till soil likely contributed to greater secondary incorporation, which was indicated by a higher diversity of late incorporators for both xylose (day 14) and cellulose (day 30). While the rate of xylose mineralization peaked on day 2 in both tillage regimes, we observed a smaller and less diverse group of early xylose responders in PTH soils relative to NTH. Early, dynamic growth of high *rrn* xylose incorporators in NTH microcosms fueled higher diversity of day 14 late incorporators and longer persistence of labeled xylose in the bacterial community. If C is rapidly metabolized, it may have greater opportunity to be cycled by diverse members of the community, undergoing extended *in vivo* processing by the microbial community (Liang et al., 2017; Fig 8). Microbial incorporation and transformation of C is recognized as a dominant process by which C is stabilized in soil (Lavallee et al., 2020). Thus, our results underscore the importance of boom-bust growth dynamics that perpetuate the persistence of microbial-derived carbon. Microbial growth and death are fundamental to C biochemical processing (Sokol et al., 2022), and we demonstrate that a tillage disturbance legacy is associated with lagged ruderal growth, thereby altering downstream processing.

**Figure 8.**
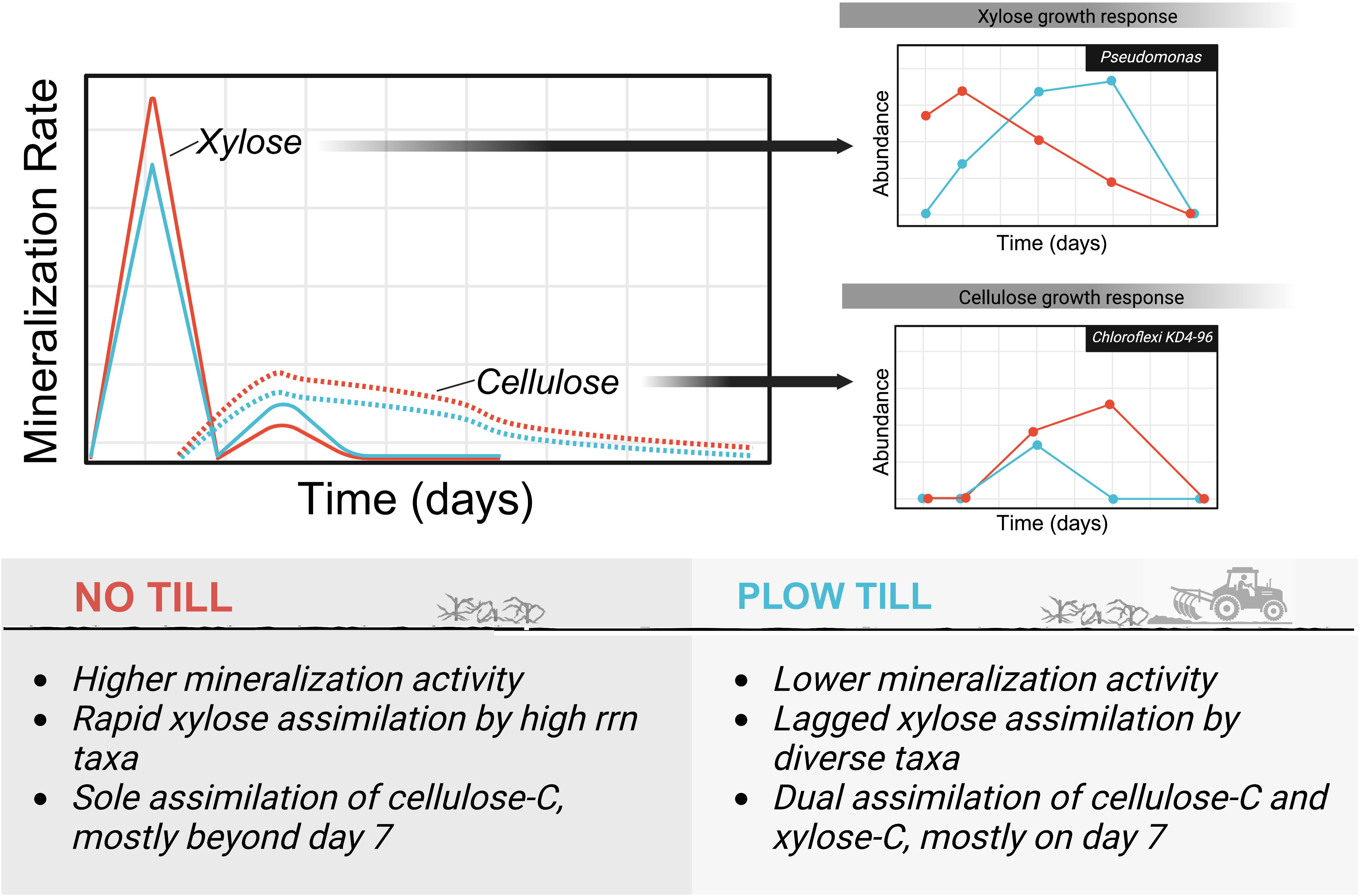

While xylose assimilation had higher latency in plow-till soils, cellulose assimilation had higher latency in no-till soils (Fig 7B). The growth dynamics of early cellulose responders were similar between the tillage regimes, indicating that the higher latency in no-till derived from a greater presence of low *rrn* incorporators on days 14 and 30 (Fig 6). Our finding that tillage regime did not alter early cellulose dynamics concurs with previous work showing that bacterial assimilation of insoluble carbon is typically not limited by growth rate, but rather by other factors such as surface colonization and enzyme diffusion (Barnett et al., 2021; Wilhelm et al., 2021). Cellulose labeling peaked between days 7 and 14 in both PTH and NTH microcosms, which coincided with peak cellulose mineralization rates (Figs 3, 6). Initially, higher species richness and phylogenetic diversity of cellulose incorporators in PTH soil was driven by a more than an 11-fold difference in the number of dual-labeled taxa on day 7 relative to NTH soil (Fig 6B). However, continued secondary incorporation and persistent growth of low *rrn* cellulolytic taxa, such as *Cellvibrio* and *Devosia* (Fig 6), resulted in a 4-fold greater median species richness of cellulose-incorporators in NTH soil by day 30 relative to PTH. Higher latency of cellulose assimilation in NTH soil suggests that minimal disturbance management resulted in longer *in vivo* processing times that retained cellulose-C in bacterial biomass for up to 30 days.

Furthermore, we did not find evidence to support a predictive role for incorporator phylogeny in explaining differences in C cycling across the two tillage regimes. Previous stable-isotope probing experiments found that incorporation of bioavailable C by soil bacteria is phylogenetically constrained, with *Actinobacteria* demonstrating consistent growth in response to glucose (Goldfarb et al., 2011; Morrissey et al., 2017). We found that incorporator phylogeny was a relatively poor predictor of C incorporation traits, and the relationship between phylogenetic distance and functional distance was especially marginal in plow-till soil (Supplemental Figure 8).

### 4.4. Life history explains incorporator carbon use but not differences in assimilation between tillage regimes

Life history strategies, including Grime’s competitor-stress tolerator-ruderal (CSR) framework (Grime, 1977), can guide predictions regarding the effects of land use on microbe-mediated C cycling (Malik et al., 2020). We predicted that a 42-year history of tillage would favor the proliferation of ruderal taxa owing to a low stress, high disturbance environment (Grime, 1977). Because both tillage regimes were managed conventionally with fertilizer input, we expected ruderals to be highly abundant in both regimes owing to pulses of high nutrient availability. We also predicted that a long-term no-till history would create a low-stress, low-disturbance environment, resulting in a higher abundance of competitors in the no-till system. Within intensively managed arable soils, we did not expect scarcity-adapted taxa to play a significant role in short-term C cycling owing to a lack of resource limitation.

To examine tillage impacts on the life history characteristics of C cycling bacterial communities, we calculated the predicted *rRNA* copy number of labeled ASVs (Stevenson and Schmidt, 2004; Wattenburger and Buckley, 2023). In agreement with previous reports, we found that *rrn* was a reliable predictor of carbon use preference (Barnett et al., 2021). Xylose, which is highly soluble, was preferentially assimilated by high *rrn* organisms, while insoluble cellulose was assimilated primarily by low *rrn* organisms. Our results confirmed our expectation that growth-adapted taxa with high *rrn* preferentially consume xylose, which is soluble and readily available for diffusive transport (Chaillou et al., 1999). Competition is higher for insoluble carbon sources such as cellulose, in which assimilation by bacteria is limited by surface colonization and extracellular metabolism (Wilhelm et al., 2021). We interpret the lower median *rrn* of cellulose incorporators relative to xylose incorporators as evidence that cellulose metabolism was predominantly conducted by competitor taxa in both tillage regimes.

We found that the median *rrn* of cellulose incorporators did not differ between tillage regimes, although xylose incorporator *rrn* was higher in the no-till legacy. Concurrent with higher *rrn,* NTH xylose incorporators also demonstrated significantly lower latency and higher maximum log2-foldchange. These results indicate that tillage reduced the dynamic growth potential of high *rrn* taxa that respond to xylose. Therefore, tillage did not alter C mineralization dynamics solely by favoring the proliferation of ruderal organisms over competitor taxa. Rather, the observed differences in xylose and cellulose mineralization were more likely driven by lagged growth of high *rrn* ruderal incorporators in PTH soil. We hypothesize that disturbance altered the successional state of microbial communities in tilled soil, contributing to lower initial abundance of active C-cycling bacteria and the observed lag in ruderal growth.

We also found that incorporator *rrn* was correlated with a number of functional responses related to growth and C assimilation dynamics (Supplemental Fig 8). As expected, *rrn* correlated with max l2fc in both tillage regimes, reinforcing the utility of *rrn* as a tool in predicting bacterial life history and dynamic growth potential (Stevenson and Schmidt, 2004). In NTH soil, *rrn* was negatively associated with degree of ^13^C assimilation, suggesting that low *rrn* incorporators received a high amount of label resulting from primary assimilation. This association was non-significant in plow-till soils (Supplemental Figure 8). Additionally, low *rrn* incorporators in NTH soil tended to exhibit higher latency of labeling, resulting in a strong negative association between *rrn* and latency that was absent in PTH soil. These findings suggest that life history strategy may be less useful in predicting bacterial carbon dynamics in highly disturbed environments when factors such as successional state can also influence growth response.

We attribute lagged growth response to xylose to legacy impacts of disturbance on C-cycling, though the cause of the lag remains to be determined. There are several possible explanations that our results do not address. For example, results from a prior SIP study indicated that ^13^C-labeled viral DNA increased rapidly within the first 3 days following C input (Barnett and Buckley, 2023). Anthropogenic inputs and land-use significantly alter bacteriophage populations, and habitat disturbance favors lysogenic viruses (Liao et al., 2022). Therefore, an increased viral load in tilled soil could explain the high latency and reduced C mineralization activity we observed in xylose incorporation in PTH microcosms due to host mortality.

Alternatively, our data may provide evidence for xylose competition in tilled soils between growth-adapted bacteria and ruderal fungi, possibly *Ascomycota* or *Basidiomycota*, whose abundance was altered by tillage in this system (Supplemental Figures 9, 10). The high latency we observed in ^13^C xylose assimilation in PTH soils implies that bacterial assimilation of xylose could have been secondary to fungal processing. A large body of research has confirmed that tillage alters the distribution, diversity, and abundance of fungi (Cho et al., 2017; Frey et al., 1999; Jansa et al., 2003; Zheng et al., 2022). A shift from filamentous fungi that grow by elongation in low disturbance systems towards yeast-like forms that grow by fission in tilled fields could alter bacterial-fungal competition dynamics which underlie the differences in C-cycling we observe with respect to tillage. We expect that, as with bacteria, disturbance regimes will differentially preference fungi with ruderal life history strategies, resulting in cross-kingdom competitive interactions that have consequences for carbon cycling. Our results, which focused on the movement of carbon through bacterial communities, nonetheless underscore the substantial impact of management practices in shaping microbial growth dynamics related to C cycling.

## 5. Conclusion

We found that long-term tillage alters bacterial growth responses with functional implications for C cycling. The most dramatic impacts of tillage were evidenced by lagged growth responses to xylose and a shorter duration of cellulose labeling among active C-cycling bacteria. Xylose incorporators in no-till soil had a higher median *rrn* and demonstrated a dynamic growth response to carbon, indicating that tillage may have impacted the life history of xylose incorporators, but not cellulose incorporators. Tillage altered the trajectory of carbon assimilation in bacterial communities, with a lower diversity of late incorporators for both substrates suggesting more streamlined processing by a narrower portion of the community. Future work should connect fungal and bacterial life histories, including competition and facilitation interactions, as drivers of C cycling in managed arable soils. Other factors that may be responsible for lagged bacterial growth, such as community assembly processes and viral load, should also be investigated for their potential to structure bacterial C dynamics in managed soil systems. Our results underscore the substantial impact of soil management practices in shaping the trajectory of carbon within microbial communities.

## 6. Acknowledgements

This work was supported in part by a USDA NIFA AFRI Education and Workforce Development Postdoctoral Fellowship (grant number 2023-67012-39838).

## 7. Author contributions

MS: Data curation, formal analysis, visualization, writing – original draft, writing – review and editing. CK: Conceptualization, investigation, data curation, methodology, writing – review and editing. DB: Conceptualization, funding acquisition, methodology, writing – review and editing.

