## Supplementary material for "Long-term tillage regime structures bacterial carbon assimilation": Figure Captions

**Figure 1. Schematic of microcosm experimental design.** Soil was collected from contrasting tillage histories and used in DNA SIP microcosms. All microcosms received the same set of carbon substrates (xylose and cellulose), but only one substrate was $^{13}$C-labeled in the enriched microcosms. Microcosm headspace sampling was used to determine mineralization activity of individual substrates and cumulative mineralization of all substrates. Destructive sampling of microcosm soil over time was used to assess changes in bacterial community structure and C assimilation dynamics.

**Figure 2. Cumulative amount of carbon (mg) mineralized for individual substrates of cellulose (A) and xylose (B) over time differed by tillage history.** Asterisks (*) indicate significant differences by land use (adjusted p-value < 0.05) as determined by post-hoc pairwise comparisons of linear mixed effects models.

**Figure 3. Carbon mineralization activity (mg hr-1) of cellulose (A) and xylose (B) over time differed by tillage history.** Asterisks (*) indicate significant differences by land use (adjusted p-value < 0.05) as determined by post-hoc pairwise comparisons of linear mixed effects models.

**Figure 4. Bacterial communities differ in their response to carbon ($^{12}$C + $^{13}$C) across tillage history and time.** Relative abundance of bacterial phyla in microcosm bulk soil over time (A) indicate a proliferation of *Proteobacteria* following C addition in both tillage regimes. Microbial community structure, including Bray-Curtis (B), unweighted UniFrac (C), and weighted UniFac (D) distances, varied according to days since C addition and tillage history. Vectors indicate the directional shift of community composition over time following C addition. The contribution of tillage and days since C addition to community structure was determined via PERMANOVA analysis in vegan (Supplemental Table 5).

**Figure 5. Long term tillage regimes result in differential C assimilation by bacterial taxa.** More xylose assimilating ASVs were detected in plow till soils, while a similar number of cellulose-assimilating ASVs were detected in both tillage regimes. While the majority of $^{13}$C-labeled taxa were present in both soil types (detection), tillage determined patterns of $^{13}$C labeling, indicating active incorporation of labeled substrates.

**Figure 6. Growth and diversity of incorporator taxa differ by tillage regime.** The top 35 most abundant incorporator ASVs from each tillage regime (A) grouped by phylogenetic relatedness and color coded by phylum (y-axis) demonstrated earlier growth responses to xylose in no till relative to plow till. Peak cellulose assimilation occurred between 7 and 14 days after C addition across both tillage histories. The median observed species richness of incorporator ASVs (B) differed with respect to tillage history, time, and substrate (Supplemental Table 6). Plow till C assimilation was largely characterized by dual labeling of ASVs by both 13C- xylose and 13C-cellulose.

**Figure 7. Predicted 16S rRNA copy number (A), latency (B), degree of labeling (C), and maximum $\log_2$-foldchange (D) of incorporator ASVs differ with respect to tillage history and carbon substrate.** Significant differences (p-value < 0.05) between incorporator ASVs from different tillage histories were determined by Wilcoxon rank-sum tests.

**Figure 8. Growth responses of incorporator taxa contribute to differences in mineralization activity observed for each carbon substrate by tillage regime, indicated by color.** Plow till soils were associated with lower mineralization rates for both xylose and cellulose. Xylose incorporators demonstrated a delayed growth response in plow till soils, with maximum abundance and labeling occurring well after peak xylose mineralization. Many cellulose incorporators in plow till soils were also dual labeled with xylose, while no till cellulose incorporators had high latency and did not assimilate xylose. Average growth curves of incorporators belonging to the *Pseudomonas* and *Chloroflexi KD4-96* genera are representative of this general trend. Because initial community composition and population size were similar between tillage regimes before carbon addition, we hypothesize that tillage altered the successional state of bacterial communities, impacting the growth responses of active C-cycling microorganisms. Our results do not exclude the possibility that tillage altered competitive interactions between bacteria and fungi, which may have also contributed to divergent mineralization activity.
