## Supplemental Material for "Long-term tillage regime structures bacterial carbon assimilation"

### Supplemental Methods

#### DNA Extraction

We extracted DNA from bulk (unfractionated) microcosms using 2 x 0.25 g of soil from all replicates of each isotope x day x soil combinations (112 samples). We used a modified Griffiths phenol-chloroform extraction procedure (Griffiths et al., 2000), in which cells were first lysed by 1 minute of bead-beating at 5.5 $m\ s^{-1}$ in 2 mL lysis tubes filled with 0.5 g of 0.1 mm silica/zirconia beads, 0.5 mL extraction buffer (240 mM phosphate buffer with 0.5% N-lauryl sarcosyl), and 0.5 mL of phenol-chloroform-isoamyl alcohol (25:24:1). After lysis, 85 µL of NaCl (5 M) and 60 µL of a mixture including hexadecyltriammonium bromide (CTAB, 10%) and 0.7 M NaCL were added to the tube. The tube was then vortexed, chilled on ice for 1 minute, and centrifuged at 16,000 x g for 5 minutes at 4°C. The top aqueous layer was transferred to a new tube and placed on ice. The pellet was re-extracted following a similar procedure, with the aqueous layer again removed and combined with the first aqueous layer. These combined layers were washed with 1 mL chloroform : isoamyl alcohol (24:1) and DNA was precipitated with 2 volumes of polyethylene glycol solution (30% PEG 8000, 1.6 M NaCl) at 4°C. Precipitate was collected by centrifugation at 16,000 x g for 30 minutes at 4°C, the supernatant was removed, and the pellets washed with 1 mL of 70% EtOH. Finally, dried pellets were resuspended in 50 µL TE and stored at –20°C. Following extraction, DNA extracts from unfractionated samples were cleaned using illustra™MicroSpin™ G-50 columns (GE Healthcare; Buckinghamshire, UK; 27-5330-02) and magnetic bead purification (Agencourt AMPure XP purification; Beckman Coulter; Brea, CA; A63880), according to manufacturer protocols.

DNA for isopycnic centrifugation was extracted from 4 technical replicates of 0.25 g of soil, following the phenol-chloroform procedure outlined above. A subset of microcosm biological replicates was selected for isopycnic centrifugation: replicate 4 across all destructive sampling timepoints for $^{13}C$ cellulose/$^{13}C$ xylose/$^{12}C$ controls, replicates 2 and 3 for $^{13}C$/$^{12}C$ xylose on day 3, and replicates 2 and 3 for $^{13}C$/$^{12}C$ cellulose on day 30. Technical replicates of DNA extractions were pooled and selected for a size of 4 – 14 kb with a Blue Pippen Prep machine (Sage Science, Beverly, MA) according to the manufacturer's protocol.

DNA for isopycnic centrifugation was extracted from 4 technical replicates of 0.25 g of soil, following the phenol-chloroform procedure outlined above. A subset of microcosm biological replicates was selected for isopycnic centrifugation: replicate 4 across all destructive sampling timepoints for $^{13}C$ cellulose/$^{13}C$ xylose/$^{12}C$ controls, replicates 2 and 3 for $^{13}C$/$^{12}C$ xylose on day 3, and replicates 2 and 3 for $^{13}C$/$^{12}C$ cellulose on day 30. Technical replicates of DNA extractions were pooled and selected for a size of 4 – 14 kb with a Blue Pippen Prep machine (Sage Science, Beverly, MA) according to the manufacturer's protocol.

**Stable isotope probing and isopycnic centrifugation**

Isopycnic centrifugation was performed for a total of 42 samples: all isotope treatment x day x tillage samples from replicate 4 (n = 26), day 3 cellulose samples of both isotope and tillage treatments from replicates 2 and 3 (n=8), and day 30 cellulose treatments from replicates 2 and 3 (n=8). We prepared isopycnic gradients followed a modified protocol based on Neufeld et al. (2007), as described previously (Barnett et al., 2022, 2021). 6 µg of size-selected DNA from each set of pooled samples was added to the density gradient solution (1.69 g $ml^{-1}$) in a 4.7 mL polypropylene tube (Beckman Coulter, Brea, CA). The gradient solution was made from a concentrated stock of CsCl (1.9 g $mL^{-1}$) and diluted in a buffer solution containing 15mM Tris-HCl, 15 Mm EDTA, and 15 mM KCl to reach the target density of 1.69 g $mL^{-1}$. Sample tubes were centrifuged at 55,000 rpm for >66 hours at 20 °C on an Optima MAX-E ultracentrifuge (Beckman Coulter; Brea, CA) with a TLA-1 10 fixed-angle rotor.

Following centrifugation, 100 µl density DNA fractions were collected from the bottoms of the polypropylene tubes using syringe pump-mediated water displacement at a rate of 15 µl $s^{-1}$ (Manefield et al., 2002). Fractions were collected in a deep-well 96-well plate (Corning, Tewksbury, MA). Immediately after each fraction was collected, we measured its refractive index ($R_i$) using a Reichart AR200 refractometer . The $R_i$ of each fraction was corrected by subtracting the $R_i$ of the gradient buffer and water ($R_{i\ corrected} = R_{i\ observed} – R_{i\ buffer} – R_{i\ water}$) and used to calculate the buoyant density of the DNA gradient:

$$\text{Density (g mL}^{-1}\text{)} = a\ R_{i\ corrected} - b$$

In the above equation, a and b are coefficient values of 10.9276 and 13.593, respectively, for CsCl at 20°C (Birnie, 1978). Fractions in the range of 1.673 -1.774 g $mL^{-1}$ were chosen for sequencing. This density range represents DNA segments that range in GC content from 13.5 - 80% plus an additional 0.036 g $mL^{-1}$ for $^{13}C$ labeling. An average number of 23 fractions per gradient tube were used, resulting the preparation of 979 fractions for sequencing.

Collected fractions intended for sequencing were desalted using the Agencourt AMPure XP purification kit according to the manufacturer's protocol (Beckman Counter, Brea, CA). Finally, we quantified purified DNA using the Quant-IT PicoGreen dsDNA assay (Life Technologies, Grand Island, NY). Fluorescence was measured using a FilterMax F5 plate reader (Molecular Devices, Sunnyvale, CA).

**16S rRNA library preparation and sequencing**

We performed amplicon sequencing of the v4 region of 16S rRNA gene across all unfractionated (n=112) and fractionated (n=979) samples. After fractionation and desalting, DNA SIP fractions within the density range of 1.673 - 1.774 g/mL were sequenced. This density range represents DNA segments that range in GC content from 13.5-80% plus an additional 0.036 g/mL for $^{13}C$ labelling. An average number of 23 fractions per gradient column were used, resulting in the preparation of 979 fractions for sequencing.

The V4 hypervariable region of the 16S rRNA gene was targeted with the 515f / 806r primer set developed by Kozich et al. (2013). Identical protocols were followed for 16S rRNA library preparation for fractionated and unfractionated samples, except for the addition of 1.25 ul Bovine Serum albumin (BNSA, New England Biolabs) to the PRC reactions of unfractionated samples. The volume of PCR reactions was 25 ul, consisting of 12.5 ul Q5 High Fidelity Hot Start PCR Mastermix (New England Biolabs), 2.5 ul combined forward and reverse barded primer at 10 uM, 5 ng template DNA, and 0.625 ul Picogreen reagent (Life Technologies, Grand Island, NY). Picogreen was added to enable visualization of reaction process on a qPCR machine. Triplicate PCR samples were normalized using a SequalPrep Normalization kit (Invitrogen), pooled, and concentrated to 5 ng/ul. Amplicon libraries were size-selected at 400-600 bp via gel excision and extracted using Wizard SV Gel and PCR Clean-Up kit (Promega). Pooled amplicon libraries were submitted for sequencing at the Cornell Core Facility in Ithaca, NY. Samples were run on an Illumina MiSeq using V2 chemistry with 2 x 250 bp read length.

### Supplemental Results

**Supplemental Table 1. Soil characteristics of the long-term tillage experiment in Chazy, NY.** All measurements, except moisture, were taken in September 2014. Moisture was averaged over 11 sampling timepoints from July 2014 to November 2015.

| Tillage | %C | %N | C:N | pH | % Moisture | DNA Yield (ng/ul) |
|---|---|---|---|---|---|---|
| **No-till** | 2.13 ± 0.39 | 0.18 ± 0.04 | 11.58 ± 0.32 | 6.89 ± 0.93 | 16.5 ± 2.47 | 56.0 ± 8.99 |
| **Till** | 1.50 ± 0.26 | 0.10 ± 0.02 | 14.36 ± 1.12 | 7.74 ± 0.05 | 16.5 ± 6.59 | 75.9 ± 26.9 |

**Supplemental Table 2. Linear mixed effects model results evaluating the contribution of tillage regime, sample day, and their interaction on mineralization rates (mg $CO_2$ $hr^{-1}$) of total carbon ($^{12}CO_2$ + $^{13}CO_2$) and individual substrates ($^{13}$C-xylose or $^{13}$C-cellulose).** Field replicate was included as a random effect in all models.

| Substrate | Fixed Effect | *df* | *F*-value | *p*-value |
|---|---|---|---|---|
| **Total carbon** | Tillage | 1 | 20.6048 | <0.0001 |
| | Day | 15 | 482.8405 | <0.0001 |
| | Tillage:Day | 15 | 5.4415 | <0.0001 |
| **Xylose** | Tillage | 1 | 1.0044 | 0.319 |
| | Day | 11 | 178.4375 | <0.0001 |
| | Tillage:Day | 11 | 3.9743 | <0.001 |
| **Cellulose** | Tillage | 1 | 11.2464 | 0.001 |
| | Day | 13 | 22.8514 | <0.0001 |
| | Tillage:Day | 13 | 3.3237 | <0.001 |

**Supplemental Table 3. Average cumulative carbon mineralized (mg $CO_2$) from total carbon ($^{12}CO_2$ + $^{13}CO_2$) and individual substrates ($^{13}$C-xylose or $^{13}$C-cellulose) by experimental endpoints.** Microcosm headspace was analyzed for xylose mineralization up to day 14 and for cellulose up to day 22. Superscripts indicate significant differences ($p<0.05$) according to t-tests comparing cumulative mineralization from microcosm replicates between tillage regimes.

| Tillage | Substrate | Day | mg $CO_2$ | SD |
|---|---|---|---|---|
| No till | Total carbon | 14 | 9.03 | 1.90 |
| | | 22 | 9.87 | 1.96 |
| | Xylose | 14 | 1.89 | 0.10 |
| | Cellulose | 22 | 2.67[a] | 0.17 |
| Plow till | Total carbon | 14 | 7.93 | 1.86 |
| | | 22 | 8.61 | 1.77 |
| | Xylose | 14 | 1.87 | 0.06 |
| | Cellulose | 22 | 2.35[b] | 0.18 |

**Supplemental Table 4. Tillage regime and days since carbon addition drive differences in bacterial community evenness, but not other measures of alpha diversity.** Linear mixed models were used to determine the contribution of tillage and days since C addition to bulk microcosm alpha diversity, with replicate included as a random factor. Alpha diversity measures were normally distributed according to Shapiro-Wilkes tests and values from rarefied bulk microcosm soil (irrespective of isotopic label) were used as response variables for the models. The significance of fixed effects was determined by analysis of variance in base R.

| | **Fixed Effect** | ***df*** | ***F*-value** | ***p*-value** |
|---|---|---|---|---|
| **Shannon** | Tillage | 1 | 90.053 | 0.98 |
| | Day | 4 | 90.045 | 0.24 |
| | Tillage:Day | 4 | 90.045 | 0.44 |
| **Pielou's evenness** | Tillage | 1 | 8.4212 | <0.01 |
| | Day | 4 | 3.7490 | <0.01 |
| | Tillage:Day | 4 | 0.8983 | 0.47 |
| **Inverse Simpson** | Tillage | 1 | 1.4762 | 0.23 |
| | Day | 4 | 2.0927 | 0.09 |
| | Tillage:Day | 4 | 1.1512 | 0.34 |
| **Richness** | Tillage | 1 | 0.6539 | 0.42 |
| | Day | 4 | 1.6012 | 0.18 |
| | Tillage:day | 4 | 0.8880 | 0.47 |

**Supplemental Table 5. Tillage regime and days since carbon addition contribute to variation in bacterial community structure.** PERMANOVA analyses were conducted using *adonis2* in vegan (Oksanen et al., 2022).

| | **Fixed Effect** | ***df*** | $R^2$ | ***p*-value** |
|---|---|---|---|---|
| **Bray-Curtis** | Tillage | 1 | 0.06 | <0.001 |
| | Day | 5 | 0.14 | <0.001 |
| | Substrate:Day | 5 | 0.04 | 0.17 |
| **Weighted UniFrac** | Tillage | 1 | 0.07 | <0.001 |
| | Day | 5 | 0.27 | <0.001 |
| | Substrate:Day | 5 | 0.04 | 0.02 |
| **Unweighted UniFrac** | Tillage | 1 | 0.04 | <0.001 |
| | Day | 5 | 0.11 | <0.001 |
| | Substrate:Day | 5 | 0.05 | 0.05 |

**Supplemental Table 6. Incorporator diversity varies with respect to tillage regime, days since C addition, and their interaction.** Incorporator species richness and phylogenetic diversity demonstrated non-normal distributions that were not corrected by log or square-root transformations. We therefore modeled the effects of tillage regime and days since C addition on incorporator diversity with generalized linear mixed models that included microcosm replicate as a random effect. Models were fit with *glmer* from the lme4 package (Bates et al., 2015). Models for species richness were fit with a Poisson distribution (family = poisson) while models for phylogenetic diversity were fit with a gamma distribution (family = gamma). The significance of fixed effects was determined with type III Wald Chi-square tests using *Anova* from the car package (Fox and Weisberg, 2019).

| | **Substrate** | **Fixed Effect** | ***df*** | **Chisq** | ***p*-value** |
|---|---|---|---|---|---|
| **Species richness** | Xylose | Tillage | 1 | 13.374 | <0.001 |
| | | Day | 3 | 271.764 | <0.001 |
| | | Tillage:Day | 3 | 176.585 | <0.001 |
| | Cellulose | Tillage | 1 | 5.3114 | 0.02 |
| | | Day | 3 | 150.9510 | <0.001 |
| | | Tillage:Day | 3 | 15.6767 | <0.001 |
| **Phylogenetic diversity** | Xylose | Tillage | 1 | 21.168 | <0.001 |
| | | Day | 3 | 671.424 | <0.001 |
| | | Tillage:Day | 3 | 405.437 | <0.001 |
| | Cellulose | Tillage | 1 | 6.5313 | 0.01 |
| | | Day | 3 | 147.3154 | <0.001 |
| | | Tillage:Day | 3 | 37.7370 | <0.001 |

**Supplemental Figure 1. Total carbon ($^{12}C + ^{13}C$) mineralized (A) and mineralization rate (B) across tillage regimes and time since substrate addition.** Error bars represent standard deviation of mean rates and stars indicate significant post-hoc comparisons between tillage regimes derived from linear mixed effects models (Supplemental Table 2). Control microcosms did not receive carbon substrates and did not differ in mineralization rate between tillage regimes at any point during the incubation period.

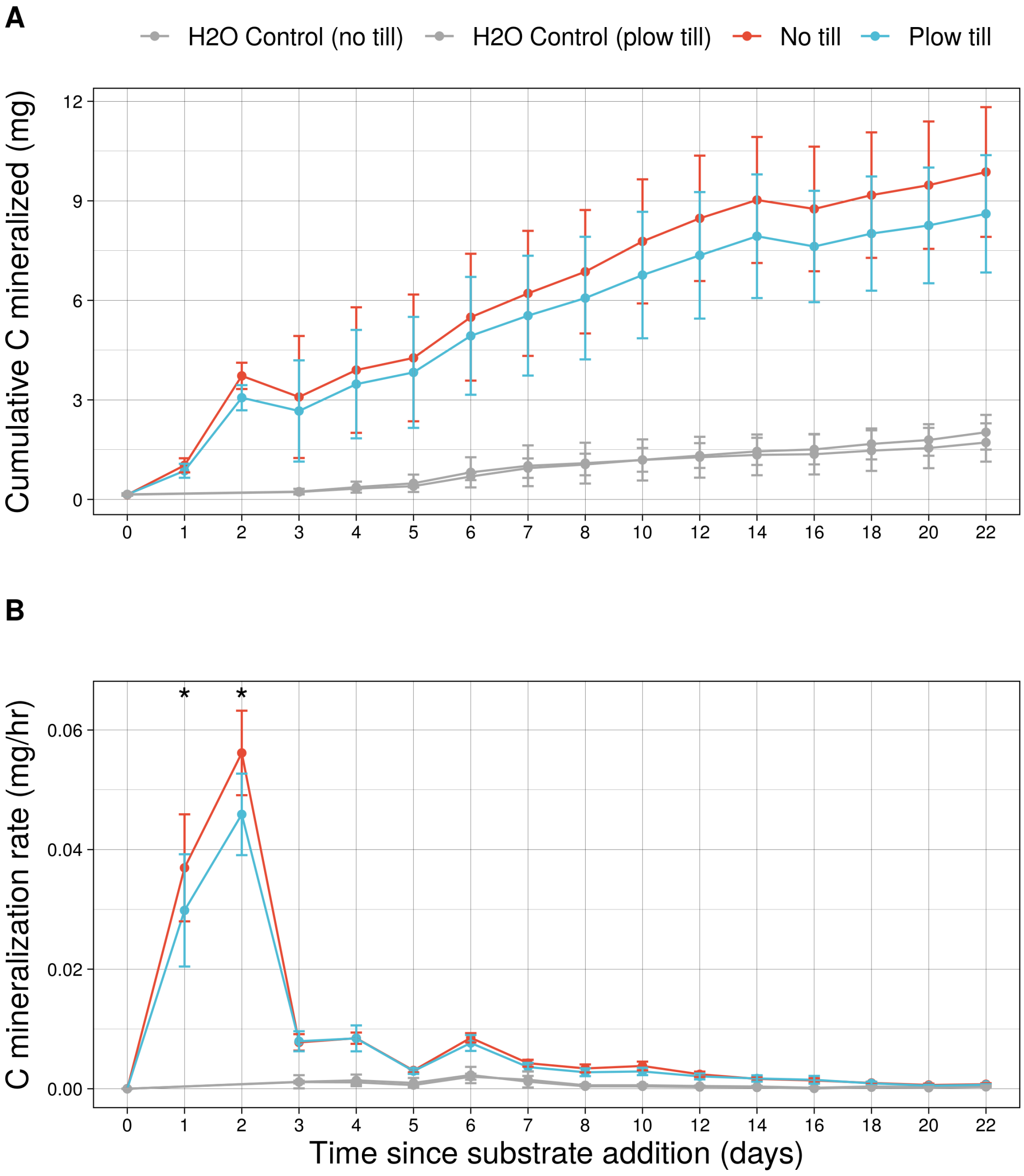

**Supplemental Figure 2. Cumulative mineralized carbon by individual substrate.** Total mineralized $^{13}C$-cellulose was determined by cumulative values up to sampling day 22, while mineralized $^{13}C$-xylose was determined by cumulative values up to day 14. Significant differences between tillage regimes were determined using t-tests within sampling day (either 14 or 22) and substrate.

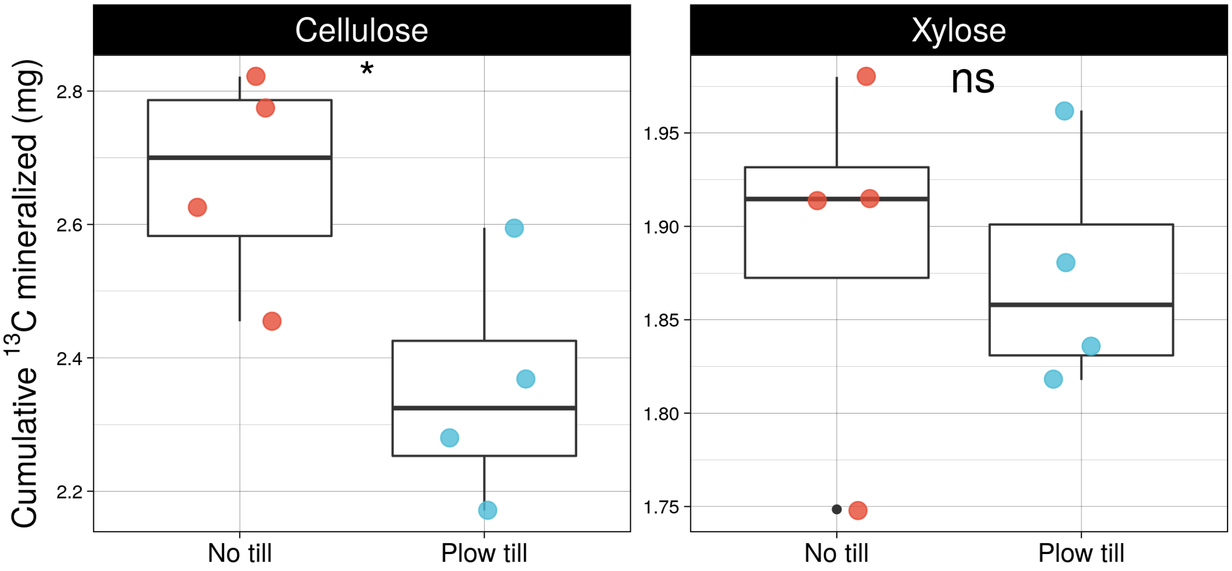

**Supplemental Figure 3. Following carbon addition, bacterial community alpha diversity tended to increase or stay the same in no till microcosms while staying the same or decreasing in plow till microcosms.** This trend was only marginally significant for the Shannon diversity index, Inverse Simpson index, and Pielou's evenness. Dashed lines indicate average diversity of water-only controls, while the shaded rectangles represent the standard deviation of controls. Significant departures from these baseline values are indicated by an asterisk. There were no significant differences by tillage regime in measures of alpha diversity prior to carbon addition.

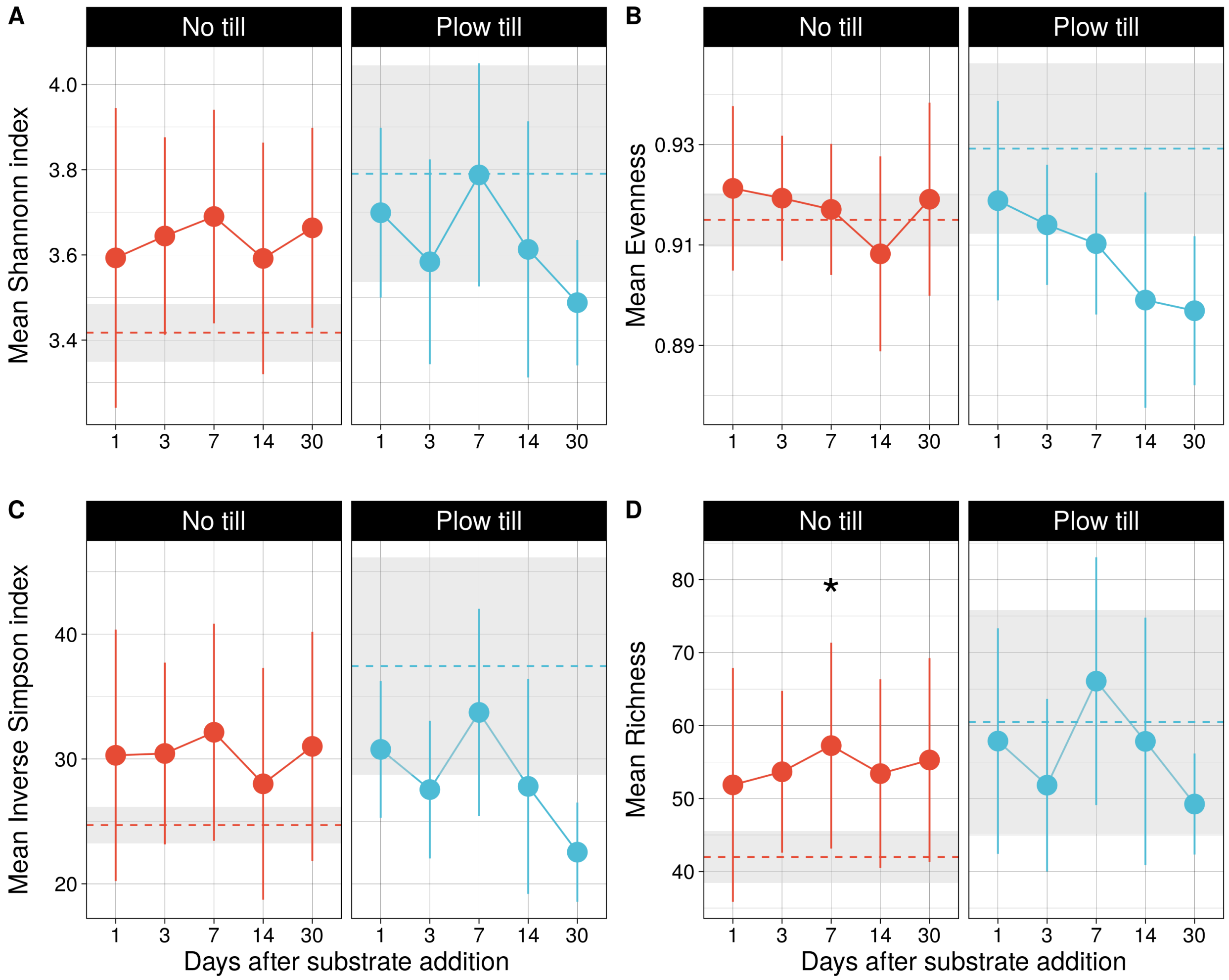

**Supplemental Figure 4. Day 30 bacterial community responses to carbon addition differ based on tillage regime.** Changes in alpha diversity measures were determined by comparing values in water-only control microcosms to microcosms that received carbon ($^{12}C$ + $^{13}C$). After 30 days, no till and plow till communities exhibited opposite shifts in alpha diversity driven by C addition. On average, C addition resulted in positive shifts in alpha diversity in no till bacterial communities while the same measures tended to decrease in plow till communities. Significant differences in the degree to which plow till and no till communities differed from controls were determined with Wilcoxon rank-sum tests by tillage regime on day 30.

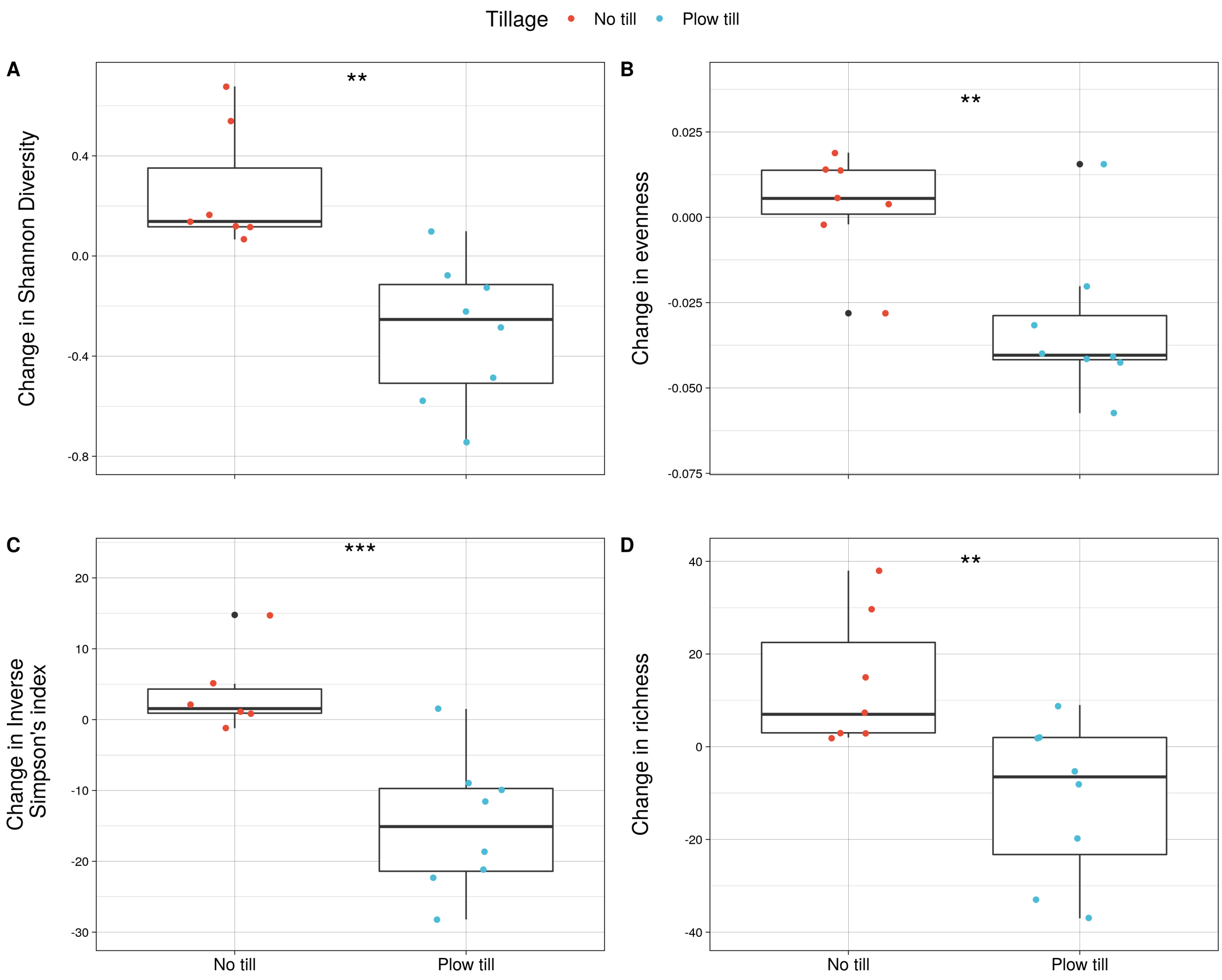

**Supplemental Figure 5. Carbon addition drives changes to microbial community structure irrespective of tillage regime.** Dashed lines represent the median distance among control microcosms that did not receive carbon inputs. Carbon addition resulted in rapid changes to community structure, with plotted data points indicating differences in beta diversity between microcosms receiving carbon and untreated controls receiving water. Significant differences in the distance between control and treated microcosms were determined via Wilcoxon rank-sum tests within tillage and sampling day. P-values were adjusted for multiple comparisons as described previously. Closed circles indicate significant differences in beta diversity compared to water-only controls.

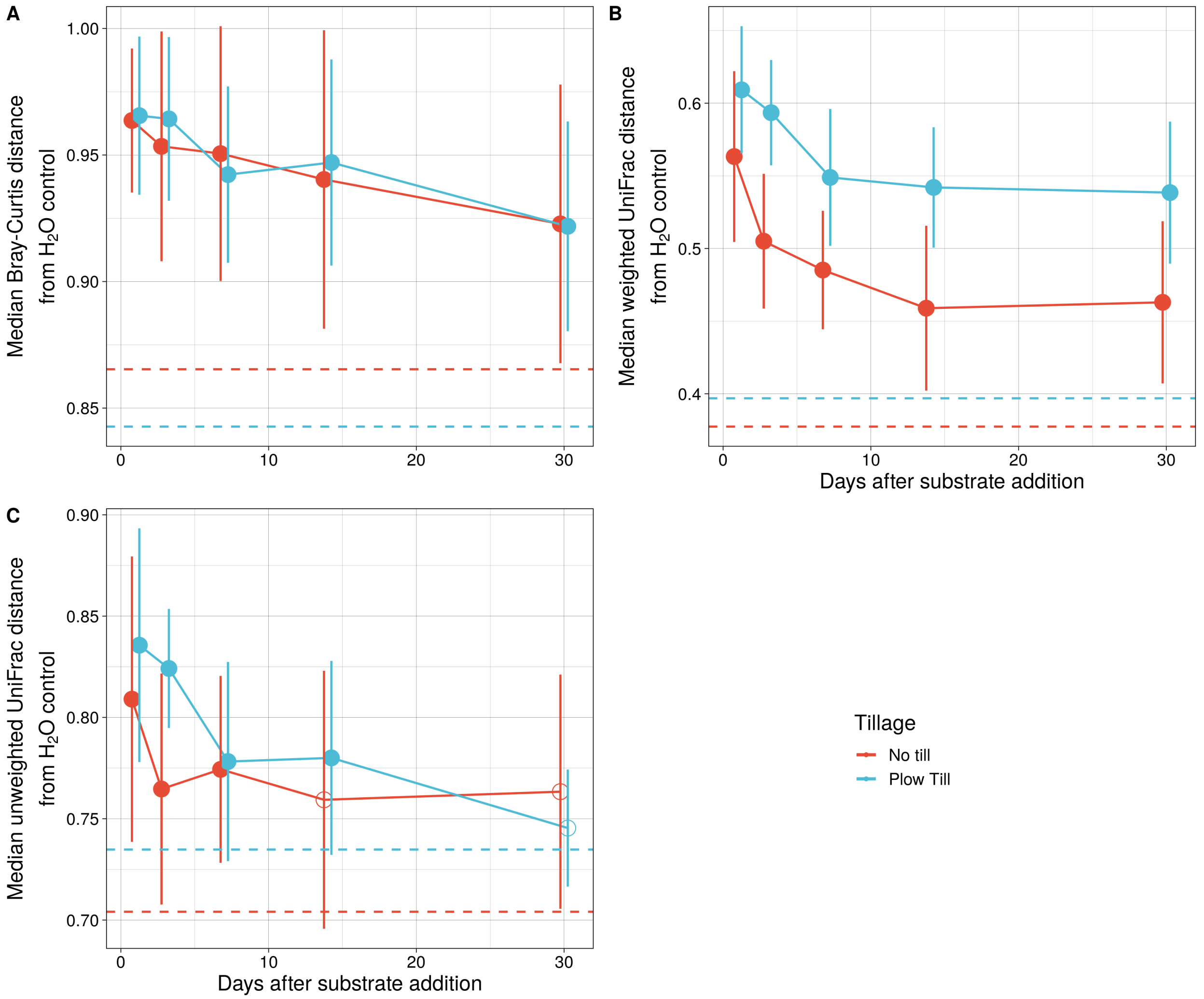

**Supplemental Figure 6. Incorporator diversity varies by labeled substrate, tillage regime, and days since C addition.** Significant differences in the phylogenetic diversity (A) and species richness (B) of incorporator taxa were determined by post-hoc tests of generalized linear mixed models (Supplemental Table 6).

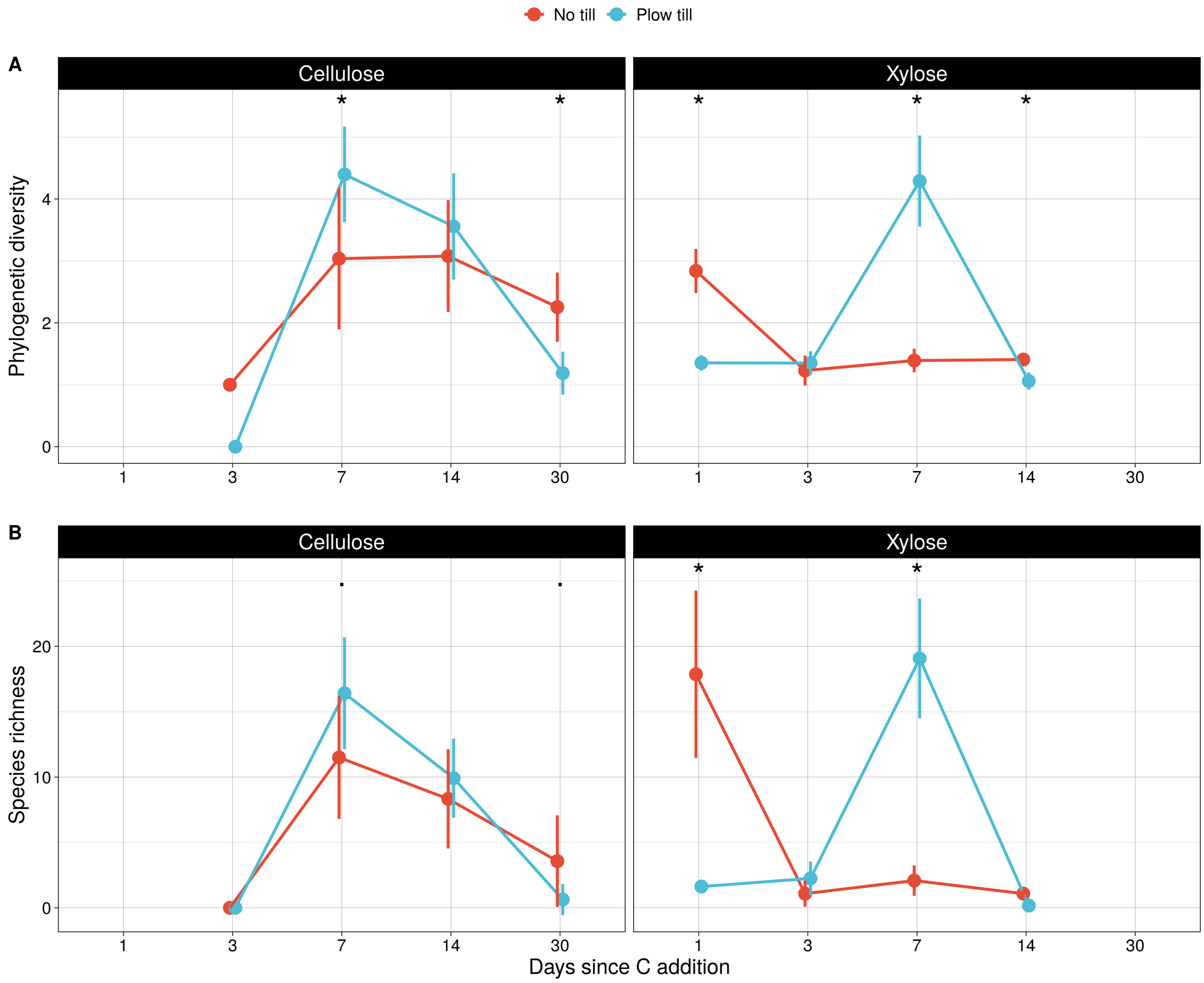

**Supplemental Figure 7. High *rrn* (> 3) and low *rrn* (< 3) incorporator taxa differ in their growth response by carbon substrate and tillage regime.** Normalized abundances were calculated as described previously by standardizing relative abundance with DNA yield and 16S rRNA copy number. The day of maximum observed abundance was determined for each incorporator ASV that was detected in rarefied microcosm soil communities. Differences in the growth by *rrn* category and tillage regime were probed with Fisher's exact test (R Core Team, 2020).

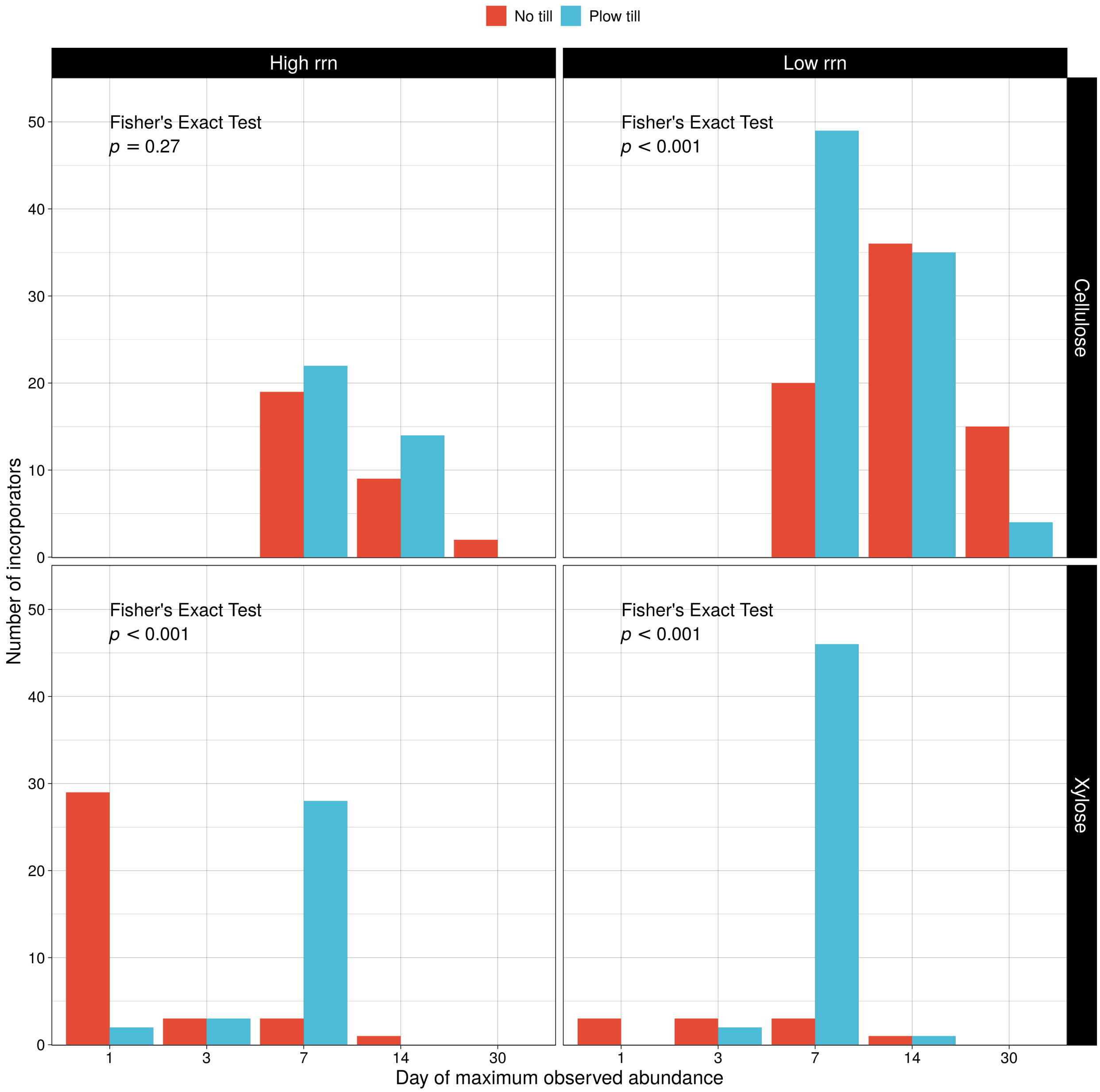

**Supplemental Figure 8. Associations between predicted 16S rRNA copy number (*rrn*) and growth responses vary among incorporators from contrasting tillage regimes.** Spearman's correlations were used to determine associations between log predicted *rrn* and growth responses to carbon. Spearman's rho and p-values are reported for significant ($p < 0.05$) correlations.

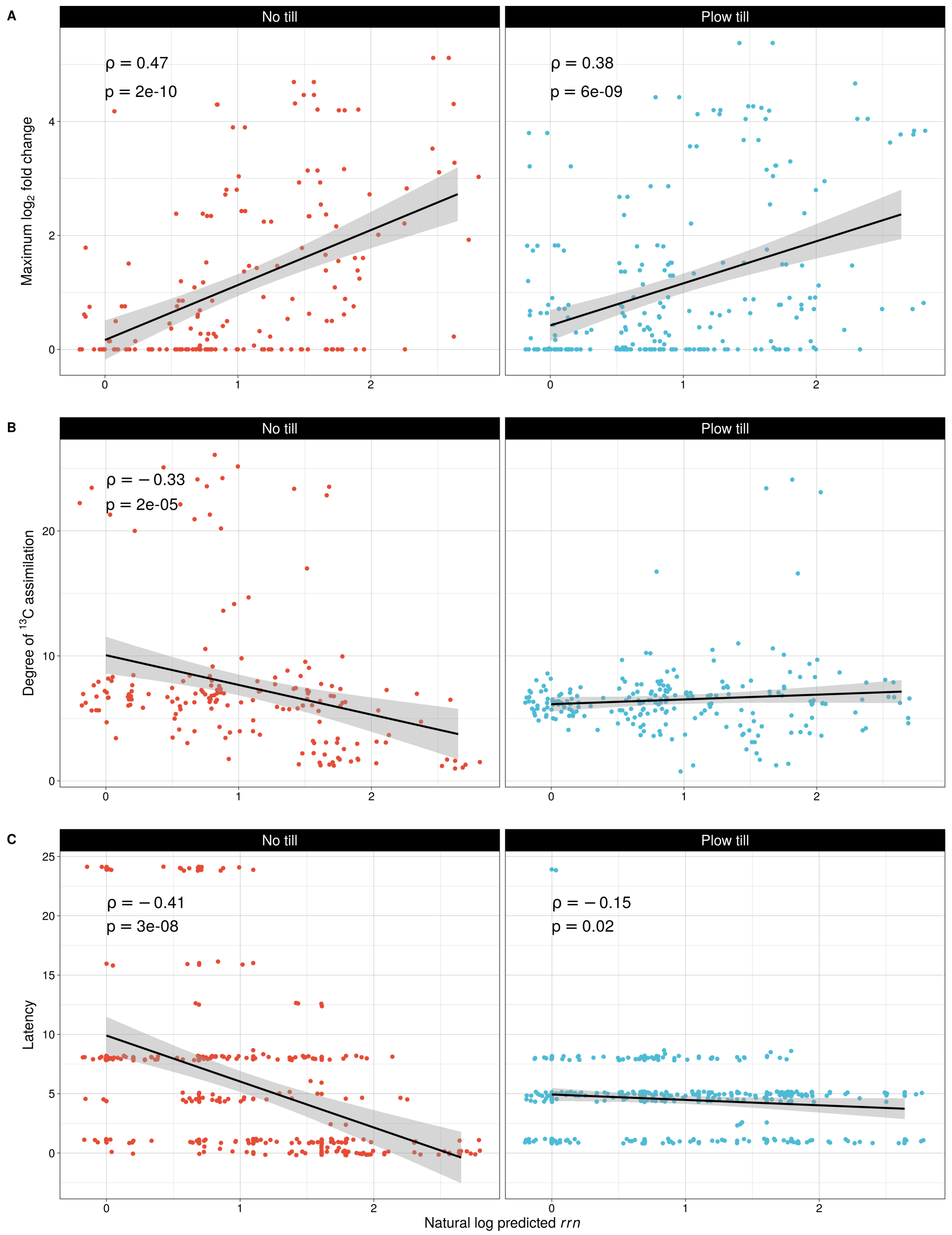

**Supplemental Figure 9. Functional characteristics related to carbon assimilation are poorly conserved among labeled taxa.** Functional distance was calculated based on incorporated substrate ($^{13}$C-cellulose or $^{13}$C-xylose) day of labeling, and degree of $^{13}$C enrichment (see methods). Dashed lines denote expected distance based on average branch length for genus, family, order, class, and phylum (left to right). Incorporator ASVs beyond the genus level exhibit substantial functional dissimilarity.

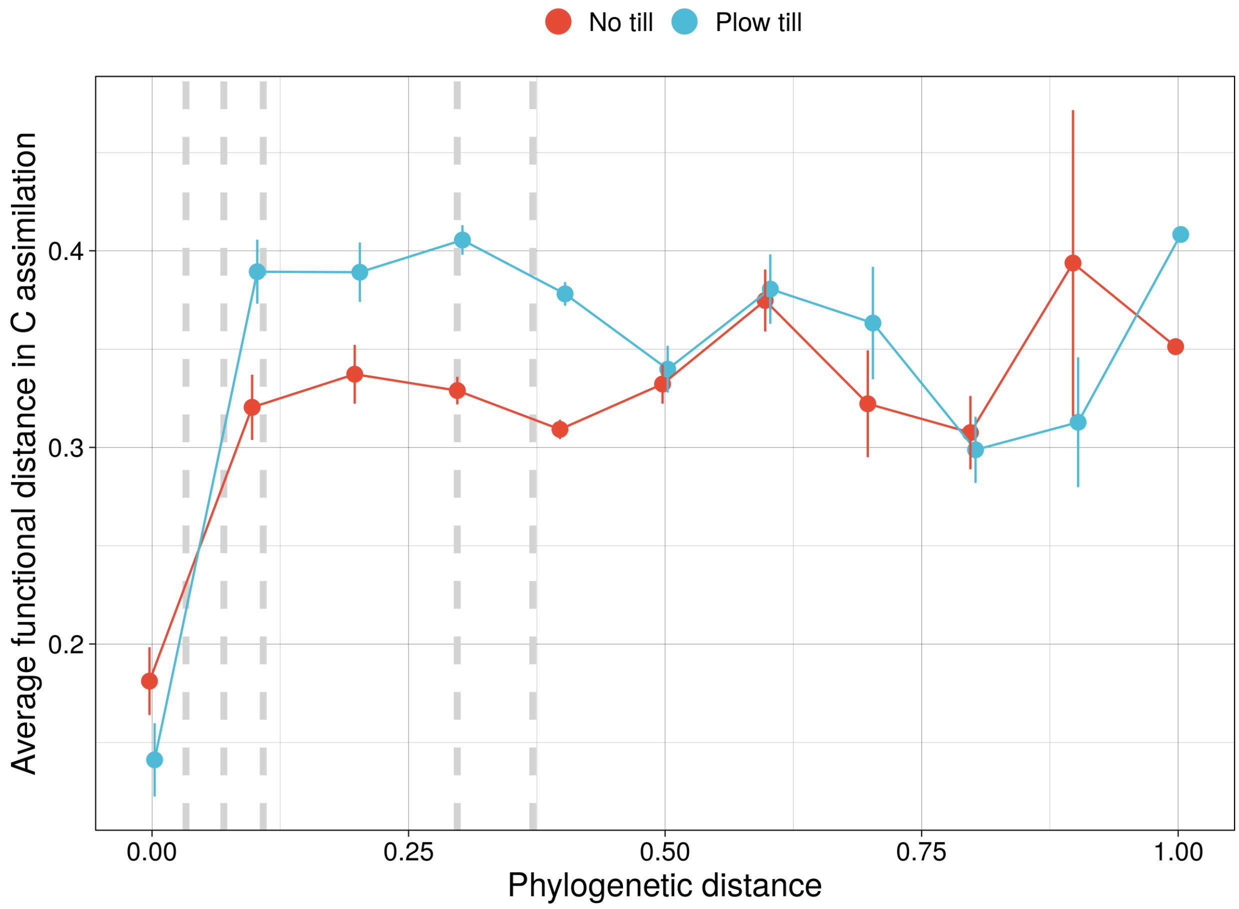

**Supplemental Figure 10. Log$_2$-fold change of fungal OTU abundance in till (positive values) vs no till (negative values) arranged by fungal order for *Ascomycetes*.** Soil samples were collected from the long-term tillage experiment at Chazy, NY between July 2014 and November 2015. Internal transcribed spacer 1 (ITS1) amplicons were sequenced at the Cornell Core Facility in Ithaca, NY from extracted genomic soil DNA using the primer set nBITSf/58A2r (Bokulich and Mills, 2013; Martin and Rygiewicz, 2005). Colors denote fungal order and significantly enriched OTUs are identified by an open dot. Figure obtained with permission from Koechli, 2016.

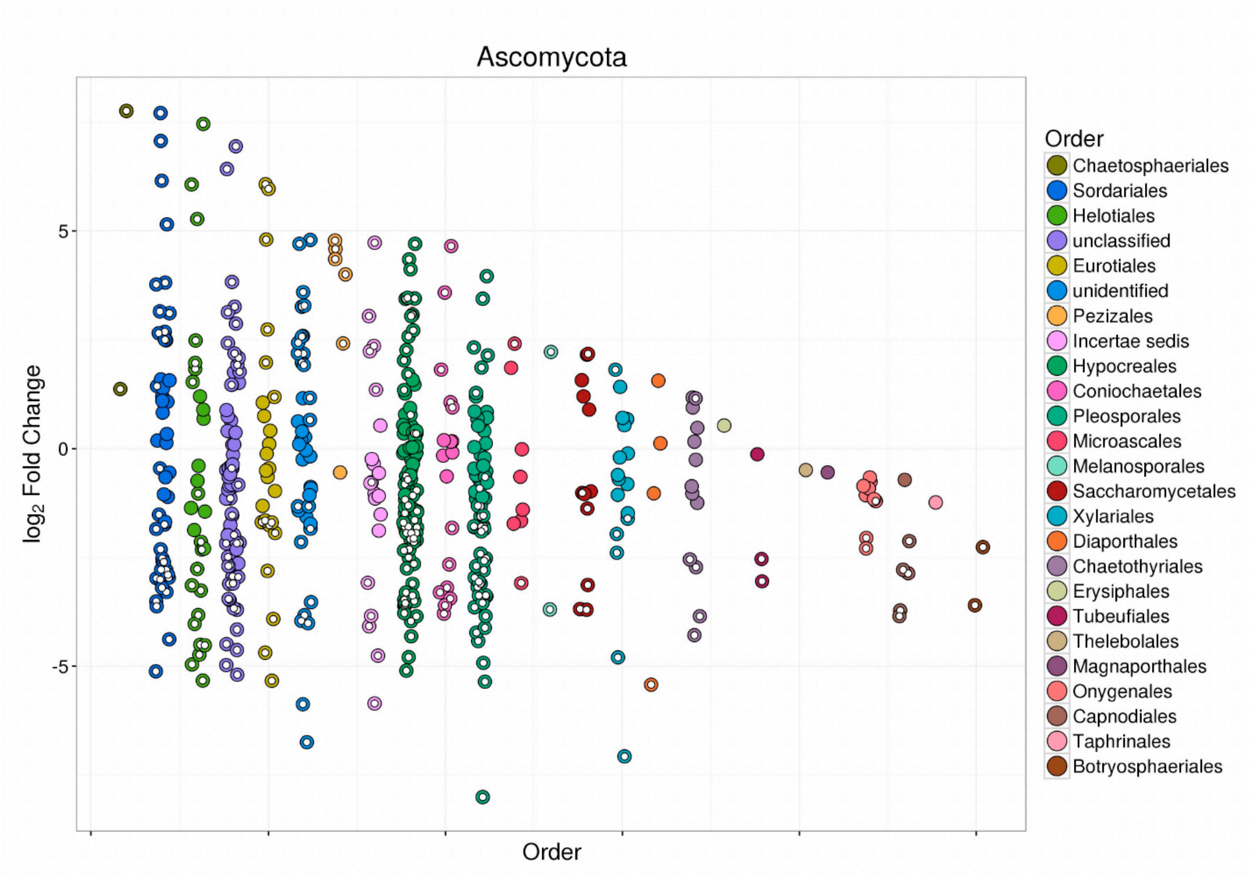

**Supplemental Figure 11. Log$_2$-fold change of fungal OTU abundance in till (positive values) vs no till (negative values) arranged by fungal order for *Basidiomycetes.*** Soil samples were collected from the long-term tillage experiment at Chazy, NY between July 2014 and November 2015. ITS1 amplicons were sequenced at the Cornell Core Facility in Ithaca, NY from extracted genomic soil DNA using the primer set nBITSf/58A2r (Bokulich and Mills, 2013; Martin and Rygiewicz, 2005). Colors denote fungal order and significantly enriched OTUs are identified by an open dot. Figure obtained with permission from Koechli, 2016.

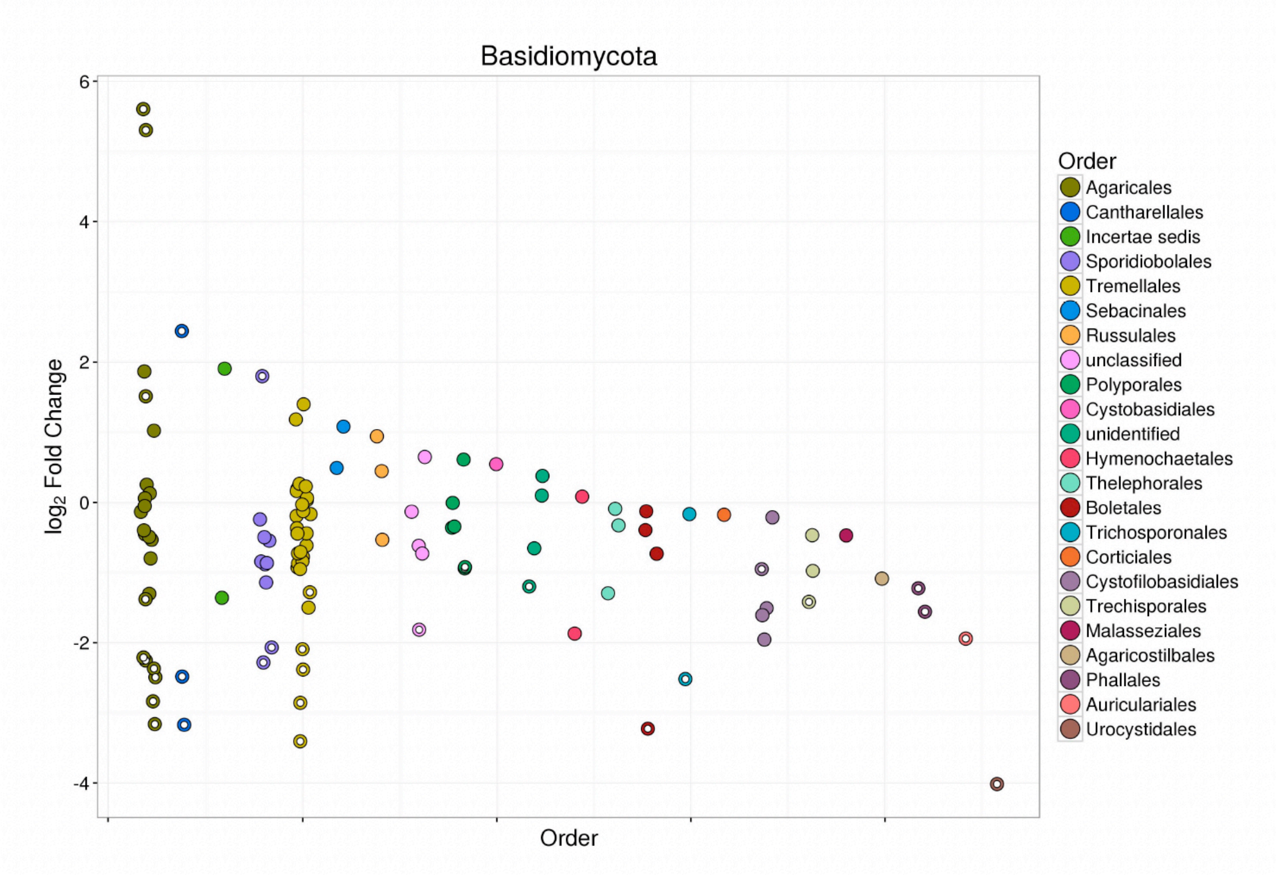
